# The Human Pancreas Cell Atlas defines a healthy reference framework for disease contextualization and translational benchmarking

**DOI:** 10.64898/2026.06.22.733853

**Authors:** Shrey Parikh, Daniel C. Strobl, Sara Jiménez, Julia L. Beckmann, Lucas Arnoldt, Eljas Roellin, Valerie Vandenbempt, Michael Sterr, Mandala Aije, Ha T.H. Vu, Rebecca Melton, Jie Liu, Fan Feng, JP Cartailler, Kyle J. Gaulton, Stephen C.J. Parker, Jürgen Ruland, Christian Conrad, Marcela Brissova, Françoise Carlotti, Heiko Lickert, Roland Eils, Diego Balboa, Malte D. Luecken, Fabian J. Theis

## Abstract

A central challenge in single-cell biology is distinguishing disease-associated remodeling from normal cellular heterogeneity. Addressing this challenge requires healthy reference frameworks that capture cellular diversity across individuals, technologies, and biological contexts. Here we present the Human Pancreas Cell Atlas (HPCA), a reference atlas of the healthy human pancreas integrating 815,126 single-cell and single-nucleus transcriptomes from 109 donors across 12 studies, diverse technologies, and demographics. Using benchmarked integration and community-driven annotations, HPCA defines 94 cell types and transcriptional states spanning endocrine, exocrine, immune, and stromal compartments. The atlas identifies rare endocrine populations, including a putative, spatially supported polyhormonal alpha–beta–delta state, and provides a unified framework for interpreting pancreatic cellular variation across diverse biological and demographic covariates. Projection of disease and model-system datasets onto HPCA contextualized endocrine and epithelial remodeling relative to healthy pancreatic states. Diabetes-associated endocrine cells remained embedded within the healthy endocrine state space while exhibiting disease-specific changes, as supported by spatial and eQTL concordance analyses. Integration with a pancreatic ductal adenocarcinoma atlas resolved injury-associated and malignant epithelial ecosystem regions across donors. Finally, the HPCA enables quantitative benchmarking of murine diabetes models and stem-cell-derived islets against human pancreatic reference states. Together, the HPCA establishes a healthy transcriptional coordinate system for interpreting disease-associated pathophysiology, experimental perturbation, and regenerative fidelity, illustrating how reference atlases can function as analytical frameworks rather than static cell catalogs.

## Introduction

In humans, the pancreas is a vital organ central to digestive and endocrine homeostasis, with exocrine and endocrine compartments implicated in some of the most prevalent and lethal diseases globally^1–5^. Single-cell RNA-sequencing studies have uncovered important insights into pancreatic cellular heterogeneity^6^, developmental processes^7^, and disease-associated cell states^8,9^. However,transcriptomic data from healthy pancreas remain fragmented and sparse, often collected within disease-focused studies, while differences in sequencing modalities, tissue processing, annotation strategies, and regional sampling have hindered the establishment of a unified reference framework for the field.

Integrated single-cell atlases have emerged as essential resources for resolving cellular variation across tissues and enabling consistent interpretation of new datasets^10–12^. Large-scale initiatives such as the Human Islet Research Network (HIRN)^13^ with its data generation program Human Pancreas Analysis Program (HPAP)^14^, together with efforts including the ESPACE Consortium^7^, as well as individual investigator-driven studies^6,8,9,15–21^ enabled by programs such as the Integrated Islet Distribution Program^22^ and the Alberta Diabetes Institute IsletCore^23^, have substantially expanded and integrated pancreatic single-cell and single-nucleus datasets across healthy and diseased states. Nevertheless, cross-study comparisons remain complicated by differences in study design, tissue sampling, sequencing modality, preprocessing, quality-control thresholds and annotation strategies. These challenges are amplified in the pancreas by rapid RNA degradation driven by abundant exocrine RNases and pervasive ambient RNA contamination from highly expressed exocrine and endocrine transcripts. Thus, a harmonized healthy human pancreas reference is needed that systematically reprocesses independent studies under unified quality-control, benchmarking, integration and annotation criteria while jointly resolving endocrine and exocrine compartments across donors, anatomical regions and technologies.

Beyond cataloguing pancreatic cell identities, such a healthy baseline is essential for determining whether disease-associated cellular programs represent discrete pathological identities, shared stress responses or continuous remodeling of pre-existing healthy states. In type 1 diabetes (T1D), beta cells exposed to autoimmune attack acquire inflammatory and stress-associated programs linked to immune-endocrine cross-talk^8,9^, whereas type 2 diabetes (T2D) is associated with trajectories of metabolic exhaustion, oxidative stress, and partial dedifferentiation^24,25^. In the exocrine pancreas, injury and chronic inflammation induce epithelial plasticity, including acinar-to-ductal metaplasia, regenerative remodeling and stress-adaptive programs that partially overlap with early oncogenic transcriptional states^26^. These observations suggest that pancreatic disease organization may be captured, at least in part, as remodeling of healthy programs rather than as fully discrete pathological states. Resolving these landscapes requires a healthy reference manifold sufficiently powered across donor diversity, sequencing modalities and biological variability to distinguish pathological deviations from normal heterogeneity and to project disease trajectories onto a stable coordinate system.

Here, we present the Human Pancreas Cell Atlas (HPCA), a comprehensive integrated reference and unified transcriptomic manifold of the healthy human pancreas. HPCA integrates 815,126 single-cell and single-nucleus transcriptomes from 109 donors across 12 independent studies including the Human Cell Atlas pancreas initiative ESPACE^7^, HPAP^14^, harmonized under a unified framework for quality control, benchmarking, and annotation^6,7,10^. The atlas resolves endocrine and exocrine compartments through a hierarchical four-level annotation framework capturing major lineages, cellular subtypes, transitional populations, and stress-associated states. Using HPCA, we characterize obesity-associated chronic inflammatory programs, define healthy intercellular signaling architectures and benchmark stem-cell-derived islet differentiation protocols. We further project 1,018,479 cells from 261 donors spanning healthy controls, T1D, T2D, autoantibody-positive, pancreatic ductal adenocarcinoma (PDAC) states and human stem cell-derived islets onto the healthy manifold, alongside cross species comparison with 193,435 mouse endocrine cells from diabetes models. These analyses suggest that epithelial disease organization is structured predominantly as continuous remodeling of pre-existing healthy programs and bifurcating ecosystem trajectories rather than isolated pathological states. In total, HPCA contextualizes 2,027,040 pancreatic transcriptomes providing an unprecedented-scale reference framework for interpreting pancreatic cellular identity across health, disease, cancer, model systems, and regenerative stem-cell engineering.

## Results

### The Human Pancreas Cell Atlas

We constructed a comprehensive reference atlas of the healthy human pancreas by integrating 12 publicly available and previously unpublished single-cell and single-nucleus RNA sequencing datasets comprising 815,126 cells across 109 diverse donors^6,8,9,15–21^ (Fig. 1f-i, Supplementary table 1, 2), sequencing platforms and tissue processing protocols. Systematic benchmarking of seven integration strategies using scIB^27^ composite metrics (see Methods, Extended Data Fig. 1a) across highly variable genes selection strategies and batch covariates identified scANVI^28^ as the optimal framework on both biological structure conservation and batch correction (Extended Data Fig. 1a-d).

**Figure 1.**
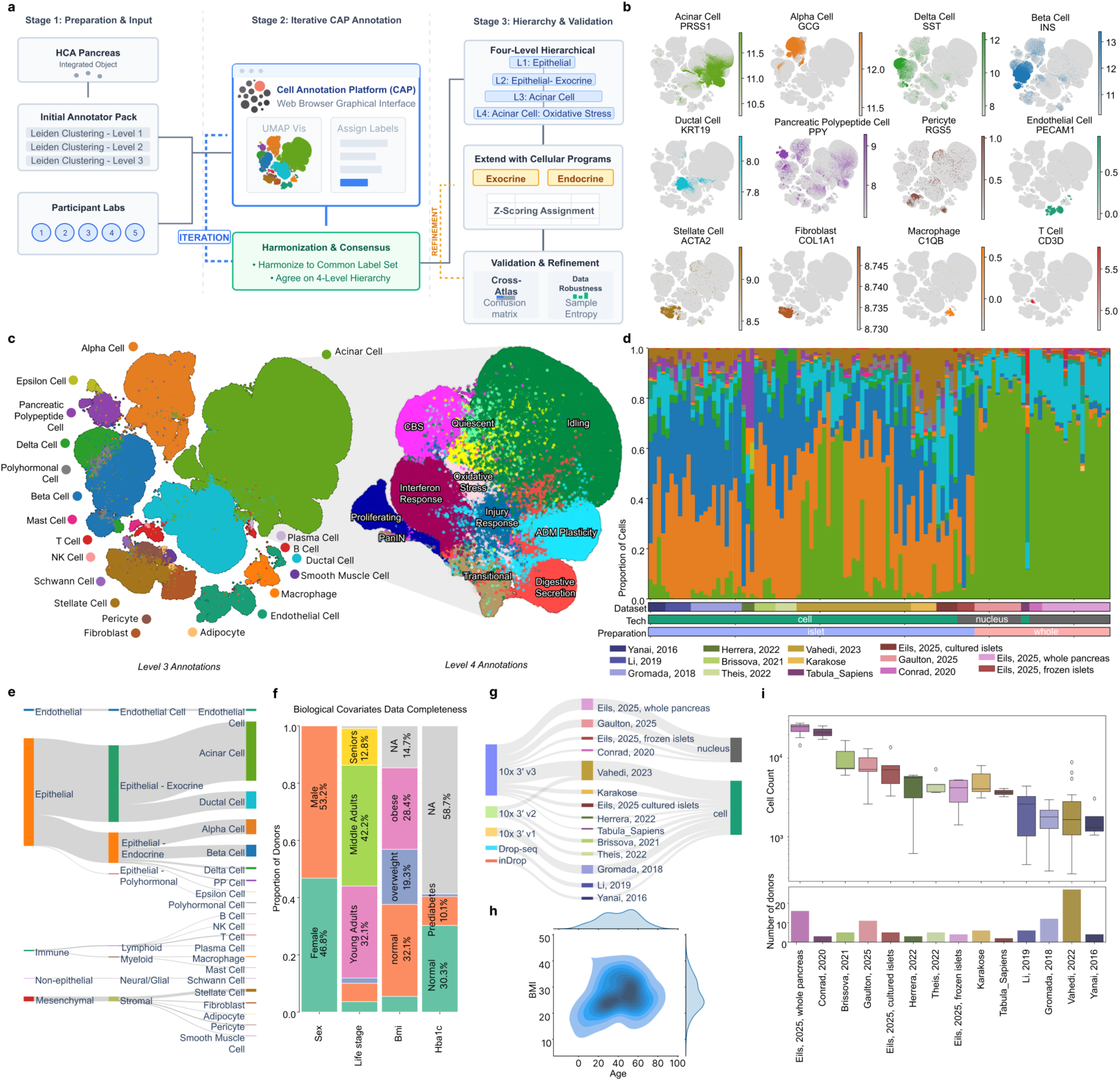
Integrated Human Pancreas Cell Atlas. **a,** Schematic overview of the annotation process. **b,** UMAP representations of the atlas coloured by expression of major lineage markers. **c,** UMAP representation of the atlas coloured by Level_3 annotation (left) and Level 4 gene program annotation of acinar cells (right) **d,** Level_3 cell-type proportions across donors and studies. The two lower colour bars indicate suspension type (single-nucleus and single-cell) and preparation method (islet isolation and whole pancreas). **e,** Hierarchical annotation up to Level_3. **f,** Metadata completeness of biological covariates Sex, Age, BMI and Hba1c. Continuous metadata fields age, bmi and Hba1c were binned for visualization purposes. **g,** Visualization of technical batch covariates across datasets. Left, sequencing platform; right, suspension type. **h,** Distribution of age and BMI across donors. **i,** Per donor cell counts and numbers of donors across datasets.

Alongside the transcriptomic data, we harmonized patient-level metadata across Human Cell Atlas (HCA) standards, including technical covariates and biological variables such as age, BMI and HbA1c (Supplementary Table 2). The atlas spans donors aged 1–80 years, BMI values of 10–45, single-cell and single-nucleus suspensions, diverse sequencing technologies and multiple sample-preparation methods (Fig. 1f-h). This breadth supports comparison of newly generated datasets across technologies, profiling strategies and donor characteristics.

To generate a robust reference, the atlas was subjected to rigorous quality control and filtering (Extended Data Fig 1 e-i, see Methods). Annotation was performed through an iterative, community-driven process using the Cell Annotation Platform (CAP) (Fig. 1a). To ensure community standards and acceptance, five laboratories across the HCA Pancreas Bionetwork independently contributed annotations to the atlas. Consensus labels were harmonized and then refined by scoring cellular programs across endocrine and exocrine compartments, yielding 94 fine-grained labels with broad concordance to available author-provided annotations (Fig. 1b, c, e; Supplementary Fig. 1a-f; see Methods). Annotation robustness was further assessed by quantifying entropy across donor, dataset and suspension type, identifying 14 clusters with limited representation diversity (Extended Data Fig. 1e, h; Supplementary Fig. 2b). Ambient RNA contamination was assessed using lineage-specific signatures, and 165,573 cells with elevated ambient RNA scores were retained but flagged in the core atlas (see Methods, Extended Data Fig. 1f, h). Because ambient RNA signatures may overlap with biologically meaningful mixed-lineage programs, these cells were retained, but all major conclusions were evaluated in the context of additional quality-control metrics (see Methods). To further improve recovery of cell states in the integrated embedding, we performed a reintegration using cell state labels to improve recovery of rare cell states (Extended Data Fig. 1j, k; see Methods).

The integrated atlas resolved the pancreas into major endocrine, exocrine, immune, endothelial and stromal compartments and established a four-level hierarchical annotation framework (Fig. 1b–e and Extended Data Fig. 2a–i,m). Broad cellular compartments and lineages were assigned at Levels 1–3, with Level 3 identities defined using canonical lineage markers, whereas higher-resolution functional and transitional states were assigned at Level 4 through a combination of gene-program scoring, thresholding (Supplementary Tables 3 and 4) and marker gene based cluster annotation. These annotations captured both established pancreatic identities and diverse programs associated with proliferation, oxidative stress, interferon response, injury and epithelial plasticity (Extended Data Fig. 2i). Within the endocrine compartment, Level 4 annotations resolved multiple alpha-, beta-, delta-and pancreatic polypeptide-cell states spanning mature, metabolic and stress-associated programs. Beta-cell states included mature-identity, secretory-granule, ion-channel, oxidative-phosphorylation, stress-responsive, proliferative and dedifferentiation-like programs, with mature identity supported by MAFA, PDX1, NKX6-1, RFX6, NEUROD1, UCN3 and IAPP^18,20,29–31^. Alpha cells similarly encompassed mature-identity, oxidative-phosphorylation, stress-responsive, proliferative and dedifferentiation-like states, with mature identity defined by ARX, MAFB, IRX2, IRX1, POU6F2, GCG, TTR and CRYBA2^18,20,32^ (Extended Data Fig. 2d). The exocrine compartment exhibited extensive Level 4 acinar and ductal heterogeneity, including digestive-secretory, quiescent, centroacinar, CFTR-transport, ionocyte-like and injury-associated states. Transitional and plasticity-associated acinar populations were characterized by SPINK1, REG1A, SOX9, KRT19, MMP7 and SPP1^33–37^, whereas injury-reactive and ionocyte-like ductal states were defined by programs containing KRT17, LGALS3, FOXI1, ASCL3 and CFTR^21,38,39^ (Extended Data Fig. 2a). The atlas further resolved pancreatic immune populations at increased granularity, including distinct B- and T-cell states (Extended Data Fig. 2b,c). Its depth and breadth enabled the consistent identification of rare and intermediate populations across donors, datasets and sequencing modalities, including interferon-responsive endocrine states, proliferative epithelial populations, transitional acinar–ductal states and polyhormonal endocrine populations. Differential-expression analysis provided complementary marker sets for each Level 4 annotation (Supplementary Table 3), establishing a high-resolution reference that distinguishes stable lineage identities from dynamic functional and remodelling states. Together, this hierarchical resolution defines a transcriptional landscape of the healthy human pancreas that is sufficiently powered to distinguish biologically meaningful cell states from technical variation and provides a reference for interpreting disease-associated remodelling against a healthy baseline.

**Extended Data Figure 1.**
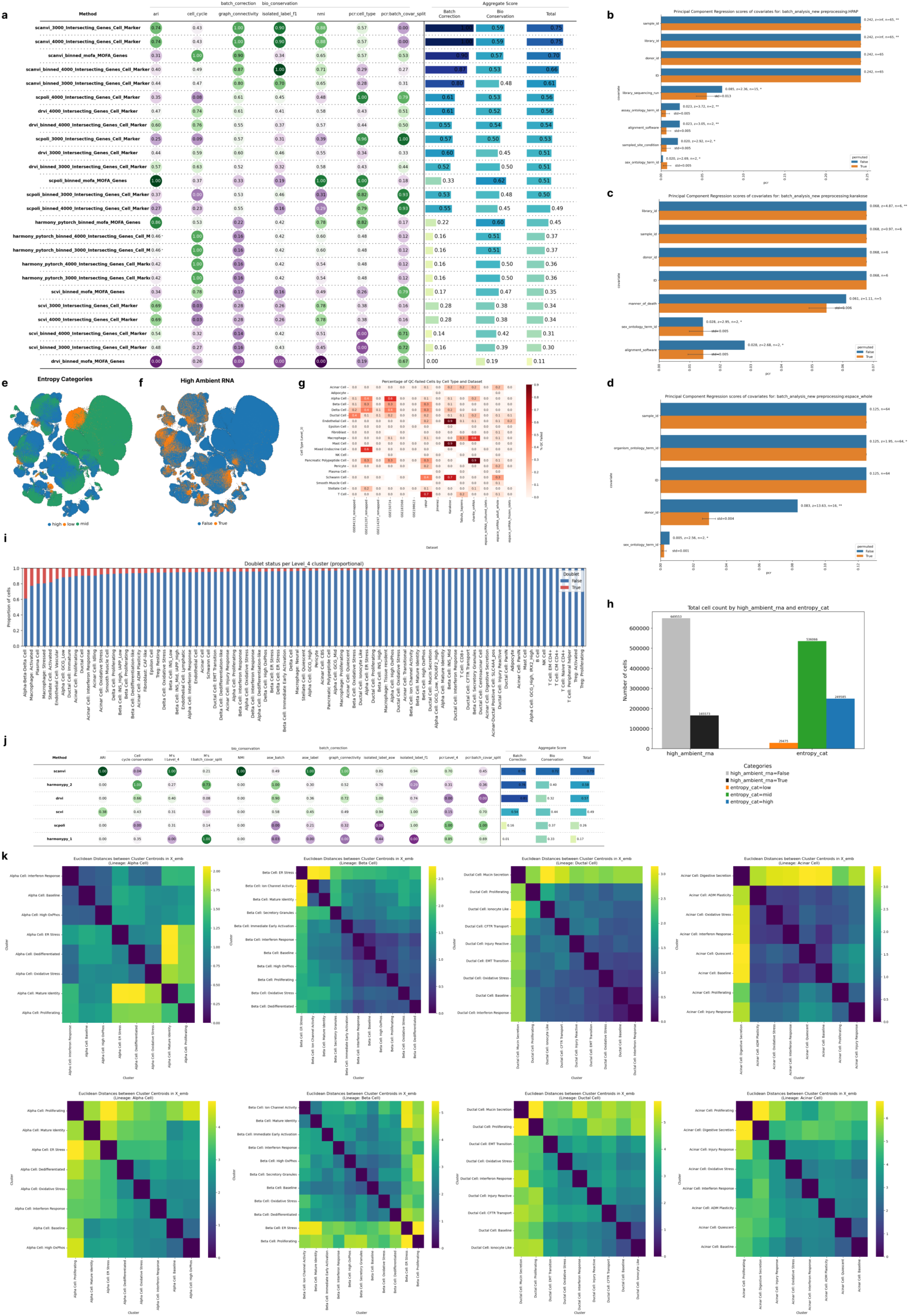
Integration benchmarking and quality control. **a,** Integration benchmarking plot showing scIB metrics across tested integration methods. **b–d,** Batch covariate analysis for datasets that were split. **e,** UMAP representation of entropy categories calculated from the mean of donor, suspension and dataset entropy. **f,** UMAP representation of ambient RNA scoring. **g,** Heatmap of final cell-level quality-control metrics per lineage. **h,** Counts of cells scored as high ambient/low entropy. **i,** Predicted doublet ratios per Level_4 label. **j,** Heatmaps of Euclidean distances in embedding space between cellular programs before (upper row) and after (lower row) integration refinement.

**Supplementary Figure 1.**
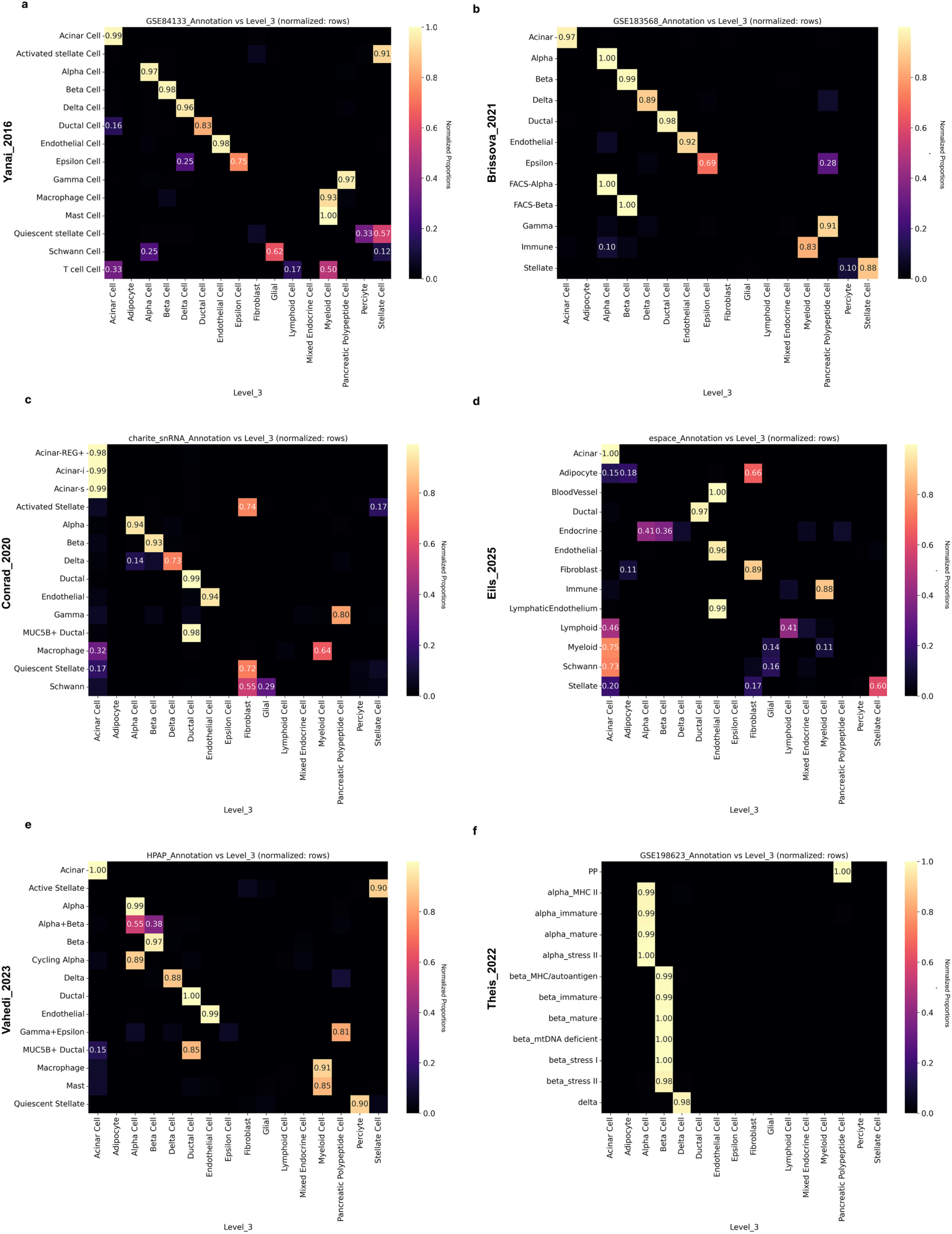
Concordance between original study annotations and HPCA Level-3 annotations. **a–f,** Row-normalized confusion matrices comparing original author-provided annotations from six independent pancreas datasets with corresponding HPCA Level-3 label assignments. Values indicate the proportion of cells from each original annotation assigned to each HPCA cell type. Strong diagonal enrichment across datasets demonstrates high concordance between study-specific annotations and the harmonized HPCA annotation framework, while off-diagonal assignments highlight differences in annotation granularity and nomenclature between studies.

### The HPCA enables multimodal identification of rare endocrine populations

We next evaluated whether HPCA-derived annotations could be transferred across modalities to spatial transcriptomic data. Using the optimal transport framework TACCO^40^, we transferred labels from the dissociated reference atlas to CosMX^9^ spatial transcriptomic data (Fig. 2a,b). Although comparison with the original author annotations yielded an average accuracy of 66% across slides (Extended Data Fig. 3e), marker-gene analysis showed that HPCA-transferred labels improved population separation and marker purity relative to the original annotations, supporting higher-resolution characterization of tissue organization and rare cellular populations in intact pancreatic tissue (Extended Data Fig. 3d,e).

**Figure 2.**
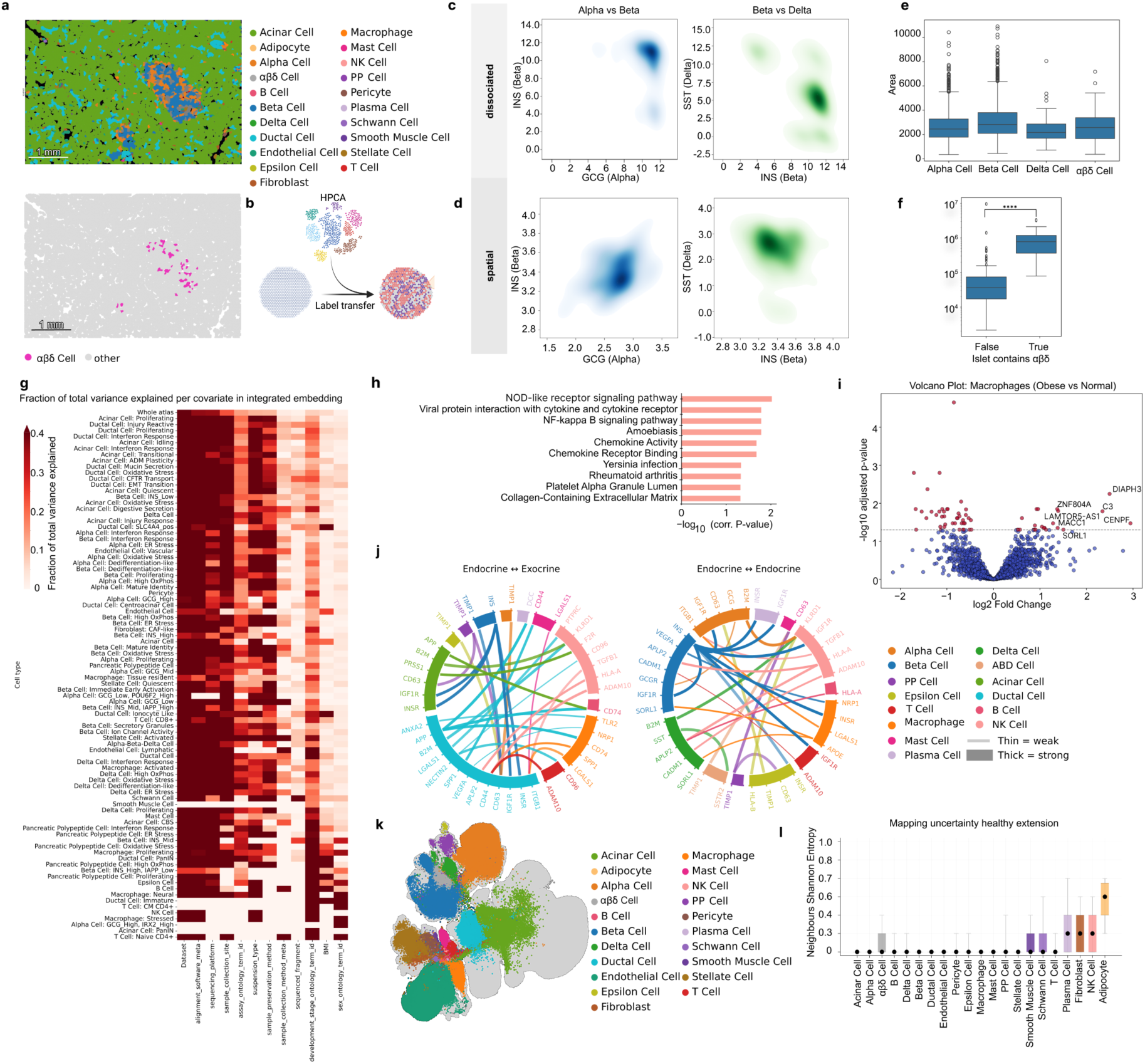
Variance association analysis, donor variability, cell–cell communication in the healthy pancreas and spatial contextualization of polyhormonal αβδ-cells. **a,** Visualization of label transfer to spatial data shown on a representative healthy pancreas slide (FOV 19, Slide 3), coloured by transferred Level_3 annotation (top). Location of polyhormonal cells within islets is highlighted (bottom). **b, s**chematic of spatial label transfer: the HPCA atlas labels were transferred to spatial data with TACCO. **c,** Density plots of marker-gene expression in dissociated data for alpha (GCG) versus beta (INS), beta versus delta (SST), and alpha versus delta markers in the αβδ cluster. Most cells in the cluster are positive for markers of all three cell types. **d,** Density plots of marker-gene expression in spatial data. **e,** Boxplot of cell areas for alpha, beta, delta and alpha–beta–delta cells, showing no size difference between Alpha, Delta and αβδ cells (Mann-Whitney U, Bonferroni corr P-Value 1) and slight size difference to beta cell (Mann-Whitney U, Bonferroni corr. P-Value 6.38e-04). **f,** Comparison of islet sizes between islets with and without αβδ cells in control slides. Islets containing triple-positive cells are larger than islets without triple-positive cells (Mann-Whitney U, P-Value 1.84e-20). **g**, Heatmap of variance contribution from technical and biological covariates across Level_4 annotations. Darker colours indicate higher variance contribution. **h,** Enriched programs of upregulated genes in acinar cells from obese donors, showing upregulation of NOD signalling, chemokine activity and extracellular matrix reorganization. **i,** Differentially expressed genes in macrophages contrasting obese (BMI>30) and normal weight donors (BMI 18-25). **j,** Cellular communication circuits in the healthy pancreas between endocrine and exocrine populations (left) and within endocrine populations (right). **k,** UMAP representation of the healthy extension mapped onto the core atlas (grey). **l,** Boxplot of mapping uncertainty stratified by Level_3 label of control cells mapped to the HPCA.

The combination of spatial mapping and high-resolution annotation further enabled investigation of 4,555 rare endocrine cells characterized by mixed alpha-, beta- and delta-cell markers. Polyhormonal alpha–beta–delta (αβδ) cells have previously been reported in stem-cell-derived islet system^41^, during development^42^ and in disease^43^, but remain poorly characterized in the healthy human pancreas. In HPCA, cultured islet datasets provided an initial discovery context for this rare state, revealing coordinated expression of alpha-, beta- and delta-associated endocrine programs, including INS, GCG, SLC30A8, IAPP, PCSK1, PCSK2, CHGA and CHGB, together with endocrine lineage regulators such as PAX6 and ISL1 and secretory-granule genes including SCG2, SCG3 and SCGN^29,44–46^. Guided by this signal, the breadth of HPCA then enabled recovery of transcriptionally similar cells from both fresh and cultured islet datasets, where fresh denotes islets profiled immediately or after brief recovery culture following isolation and cultured denotes islets maintained in vitro for extended periods before profiling, supporting the presence of this state beyond culture-enriched settings (Suppl. Fig. 2a).

To distinguish this population from mixed-lineage transcriptional profiles arising through technical artifacts, we performed multiple orthogonal quality-control analyses. Polyhormonal αβδ cells were broadly represented across donors, datasets and suspension types (Suppl. Fig. 2a,b), arguing against a donor- or dataset-specific artifact. Their mixed endocrine transcriptional profile raised the possibility of ambient RNA contamination or cell doublets. Consistent with this concern, ambient-RNA and doublet-detection methods flagged some of the αβδ cells, although most cells (approximately 60%) were predicted singlets. Additional analyses failed to support a simple technical explanation (Extended Data Fig. 2j–l; see Methods). Beta-associated scores largely overlapped with canonical beta cells (KS = 0.09), whereas alpha-associated scores showed greater separation from canonical alpha cells (KS = 0.35). Ribosomal and mitochondrial transcript fractions differed only modestly from canonical endocrine populations (KS = 0.21 and 0.26, respectively) and remained within the range observed between canonical endocrine cell types (ribo KS = 0.05–0.28; mito KS = 0.06–0.29). Furthermore, αβδ-cell frequency did not increase with dataset cell recovery (Spearman ρ = −0.13, P = 0.66), as would be expected for canonical doublets, and predicted singlets and doublets were extensively intermixed (silhouette score = 0.007). Nevertheless, because αβδ cells exhibited elevated doublet scores relative to canonical endocrine populations, downstream characterization analyses were restricted to predicted singlets, whereas all αβδ cells were retained in HPCA as a candidate endocrine state for future investigation.

Spatial label transfer provided orthogonal support for the presence of this polyhormonal transcriptional state. The label transfer identified approximately 700 αβδ cells with high confidence (> 0.7) among 384,768 spatially profiled cells. Co-expression of the canonical alpha-, beta- and delta-cell markers GCG, INS and SST was observed in both dissociated and spatial datasets (Fig. 2c,d), and marker genes identified in dissociated cells were broadly recovered in the spatial data (Extended Data Fig. 3g). We additionally observed no differences in cell size between αβδ cells and most other endocrine populations (Fig. 2e, Mann-Whitney U), arguing against segmentation artifacts. High scores for alpha-, beta- and delta-cell programs further supported the mixed-lineage transcriptional identity. Finally, islets containing αβδ cells were significantly larger than islets lacking this population (Fig. 2f), consistent with previous observations linking islet size and endocrine-cell heterogeneity^47^.

### The HPCA captures biological variation, multicellular organization and cross-cohort diversity in a generalizable reference framework

We found that both donor and technical heterogeneity contributed substantially to transcriptional variation, consistent with observations from atlases of other organs^11^. We quantified their relative contributions across pancreatic cell types and states using principal component regression for individual covariates within each annotated population^11^ (see Methods). Technical factors, including suspension type (single-cell versus single-nucleus), read alignment strategy and sample preservation method, explained a substantial fraction of transcriptional variance across many cell types (Fig. 2g). Among biological covariates, age (development_stage_ontology_id) contributed appreciably to variation in immune populations, including CD8+ T cells (31%), endothelial populations, including lymphatic endothelial cells (51%), pericytes and acinar cells, whereas BMI-associated variation was most evident in selected exocrine states, including acinar cells (20%), and proliferating macrophages (27%).

We next asked whether HPCA could resolve obesity-associated transcriptional programs. Consistent with links between obesity, chronic pancreatic inflammation, fat deposition, and exocrine dysfunction^48^, acinar cells from obese donors showed inflammatory programs characterized by increased NOD signaling^49^, chemokine activity and extracellular-matrix remodeling (Fig. 2h). Macrophages from obese donors were enriched for a proliferative state and elevated C3 expression (Fig. 2i), a marker previously associated with insulin resistance^50^. Together, these analyses demonstrate that HPCA captures biologically relevant obesity-associated variation across pancreatic cell populations.

We next explored whether HPCA could be used to characterize multicellular organization of the healthy pancreas. We inferred ligand-receptor co-expression patterns using the highly curated knowledge base and cell-cell communication framework from LIANA (Fig. 2j, see methods)^51^. HPCA captured canonical intra-islet interactions, including GCG:GCGR^52^, SST:SSTR2^53^, INS:INSR, INS:IGF1R^54,55^, as well as MHC-I-mediated immune surveillance through B2M:KLRD1^56,57^. Beyond these more established interactions, the analysis prioritized several candidate signaling axes, including VEGFA:NRP1 interactions from beta and ductal cells toward macrophages, suggesting a possible role for local VEGFA signaling beyond angiogenesis^58–60^; macrophage APOE:SORL1 interactions toward beta and delta cells, implicating lipid-handling and endosomal-trafficking pathways^61–63^; and ductal NECTIN:CD96 interactions toward T and NK cells, representing a candidate immune-checkpoint interaction with precedent in tumour immunology^64–67^. Although requiring functional validation, these interactions reveal structured communication across endocrine, exocrine and immune compartments.

A major utility of a reference atlas is its ability to support analysis of newly generated datasets, which requires the identified cellular states and embedding structure to generalize beyond the atlas-construction cohorts. To test this, we mapped 251,152 cells from 26 non-diabetic donors spanning five independent cohorts and multiple technologies^68–72^ onto the reference atlas using scArches^28^ (Suppl. Table 2, Fig. 2k,l). Level-4 labels were transferred and confidence was quantified using normalized k-nearest-neighbour Shannon entropy (see Methods). More than 90% of cells had entropy values near zero, indicating confident assignment to a single reference neighbourhood. Although cell-type compositions varied across cohorts, many differences were consistent with study-specific sampling strategies, including islet-enriched versus whole-pancreas dissociation, and may also reflect biological heterogeneity across donors and tissue regions. These findings support HPCA as a broadly applicable reference for future pancreatic studies.

Collectively, these results demonstrate that HPCA provides a stable, high-resolution coordinate system for the healthy pancreas, one that captures biologically meaningful donor variation, generalizes across independent cohorts and technologies, resolves intercellular signaling architectures, and enables high-resolution characterization of rare cell states across both dissociated and spatial modalities.

**Extended Data Figure 2.**
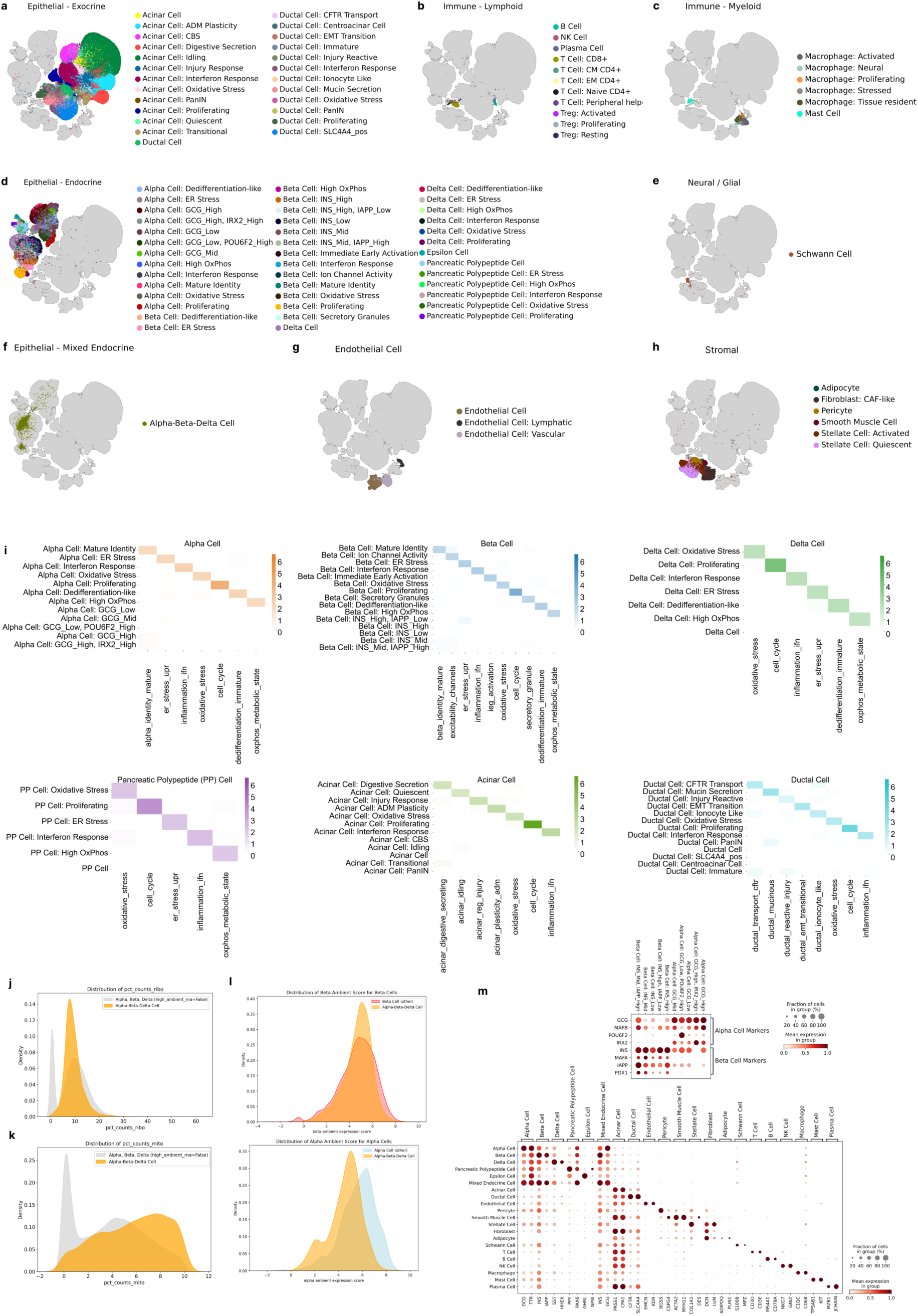
Level_4 cell-type and cell-state annotation. **a**, UMAP representation of the exocrine compartment. **b,** UMAP representation of the endocrine compartment. **c,** UMAP representation of the lymphoid lineage. **d,** UMAP representation of the myeloid lineage. **e,** UMAP representation of the endothelial lineage. **f,** UMAP representation of the neural lineage. **g,** UMAP representation of mixed endocrine cells. **h,** UMAP representation of the stromal lineage. **i,** Heatmaps Z scores of gene program scores across all endocrine and exocrine populations **j,k,** Quality-control metrics of polyhormonal cells compared with other endocrine populations. (j) Ribosomal count proportion and (k) Mitochondrial count proportions. **l,** Beta-associated ambient gene score (left) and alpha-associated ambient gene score (right). **m,** Marker-gene expression across Level_3 labels (bottom) and across Level_4 endocrine cell types defined by marker expression (top.)

**Supplementary Figure 2.**
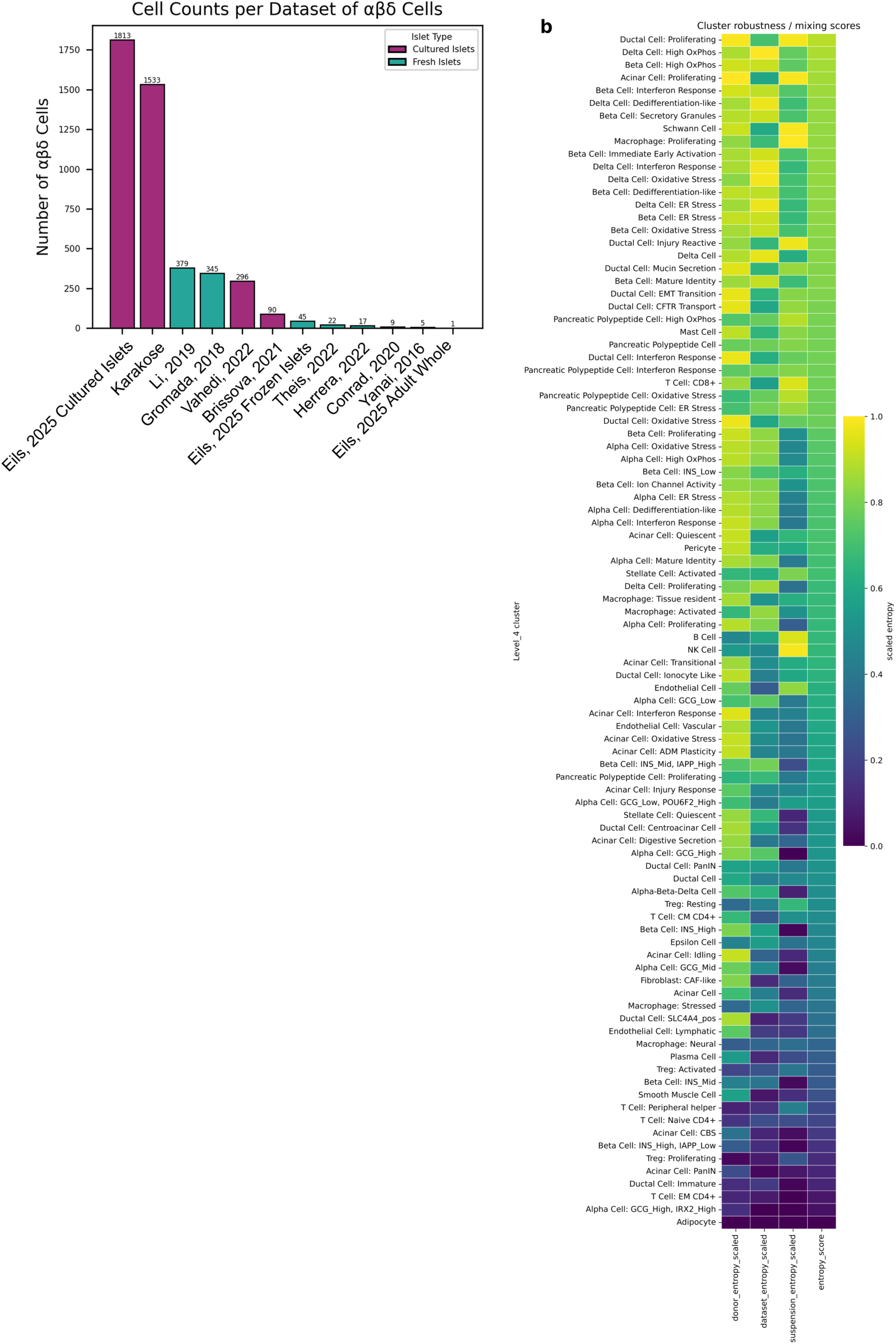
Distribution and robustness of αβδ cells in HPCA. a, Number of αβδ cells detected per dataset, colored by islet source. **b,** Scaled donor, dataset and suspension entropy, and overall entropy score, across Level-4 HPCA states. Higher values indicate broader representation across biological and technical contexts.

### Diabetes-associated endocrine states show disease-specific remodeling within healthy endocrine state space

A central question in diabetes biology is whether disease-associated endocrine states emerge as discrete pathological identities or reflect remodeling of pre-existing healthy programs. Conventional analyses typically classify cells into broad endocrine cell types, limiting the ability to determine whether disease-associated cells represent distinct identities or altered occupancy of pre-existing healthy states. To address this question, we used HPCA as a healthy endocrine reference framework onto which diabetes-associated datasets could be projected and interpreted relative to healthy endocrine state space. We mapped type 1 diabetes (T1D; 107,241 cells from 26 donors)^6,9^, type 2 diabetes (T2D; 57,635 cells from 19 donors)^6,72^ and autoantibody-positive (AAB; 101,140 cells from 21 donors)^6,9^ datasets onto the HPCA endocrine reference using scArches^28^ generating an integrated resource comprising 1,086,688 cells (Suppl. Table 2, Fig. 3a). Disease-associated endocrine cells were identified using a lineage-restricted scANVI uncertainty framework that detects cells with ambiguous assignment to healthy endocrine substates while retaining strong endocrine lineage support (see Methods). This approach identified 16,829 disease-associated endocrine cells, including 3,250 T1D, 8,774 T2D and 4,805 AAB cells. These states arose predominantly within alpha- and beta-cell compartments, with T1D-associated states enriched among alpha cells, T2D-associated states distributed across both alpha and beta cells, and AAB-associated states enriched among beta cells (Extended Data Fig. 3a).

**Figure 3.**
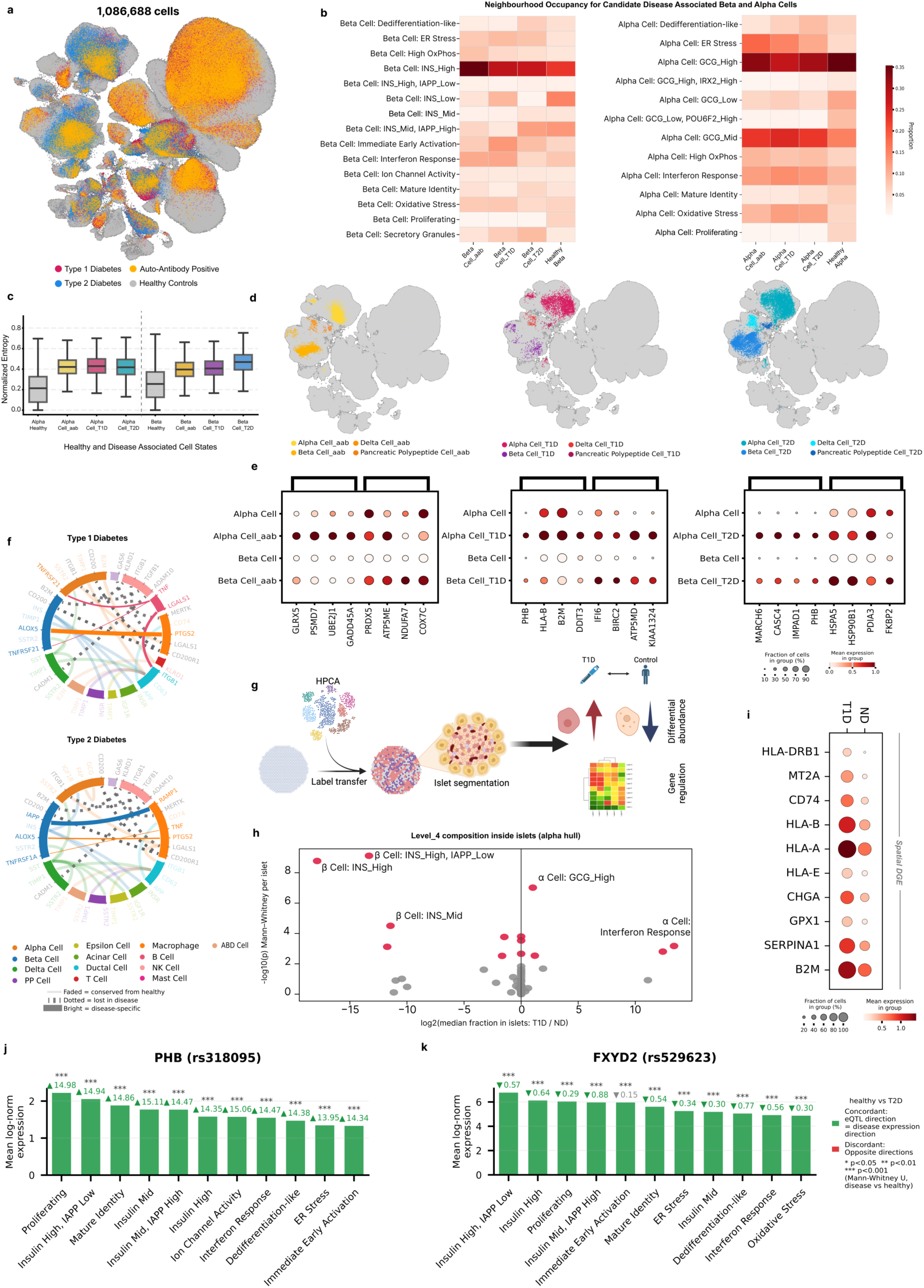
Mapping diabetes-associated endocrine states onto the HPCA endocrine reference manifold. **a,** Integrated HPCA endocrine manifold following extension with T1D, T2D and autoantibody-positive (AAB) datasets using scArches, coloured by disease condition. Healthy endocrine cells are shown in grey. **b,** Occupancy of healthy HPCA alpha- and beta-cell substates by the disease-associated endocrine cells. Heatmaps show the fraction of the disease-associated alpha and beta cells from T1D, T2D, and AAB donors assigned to each healthy HPCA endocrine substate based on k-nearest-neighbour mapping in the scANVI latent space. **c,** Weighted neighbourhood entropy of the disease-associated endocrine cells across disease conditions. Boxplots show weighted neighbourhood entropy for alpha- and beta-cell candidate disease-associated states in T1D, T2D and AAB samples. Higher values indicate broader distributions across healthy endocrine neighbourhoods and lower mapping specificity. **d,** UMAP representation of candidate disease-associated endocrine cells across the healthy HPCA for AAB, T1D and T2D donors, shown for alpha, beta, delta and pancreatic polypeptide cell compartments. **e,** Representative differentially expressed genes associated with candidate disease-associated endocrine cells in AAB, T1D and T2D donors across alpha- and beta-cell populations. Dot size indicates the fraction of cells expressing the gene and colour indicates mean normalized expression. **f,** Cell–cell communication networks inferred using LIANA for T1D (top) and T2D (bottom). Edge width corresponds to inferred interaction strength. Faded edges indicate interactions also detected in healthy controls, dotted edges indicate interactions lost in disease and highlighted edges indicate disease-enriched interactions. **g,** Schematic overview of the spatial transcriptomics analysis workflow. HPCA labels were transferred to spatial transcriptomics data, islets were segmented using alpha-cell convex hulls, and differential composition and gene-expression analyses were performed between T1D and control islets. **h,** Differential endocrine-state composition within spatially defined islets in T1D and control samples. The x axis shows log2 fold-change in median islet composition (T1D versus control) and the y axis shows −log10(P value) from two-sided Mann–Whitney tests. **i,** Differentially expressed genes in spatially resolved T1D alpha cells relative to control alpha cells. Dot size indicates the fraction of expressing cells and colour indicates mean normalized expression. **j,** Expression of PHB across HPCA beta-cell substates. Bar height indicates mean log-normalized expression within each substate. Colours indicate whether the direction of differential expression in T2D is concordant (green) or discordant (red) with the direction predicted by the colocalized T2D eQTL (rs318095, PP.H4 = 0.88). Values above bars denote T2D-versus-control log2 fold-changes. Asterisks indicate significance of differential expression. **k,** As in j, for FXYD2 (rs529623, PP.H4 = 1.00).

We next asked whether the disease-associated cells localize to specific healthy endocrine neighbourhoods or distribute broadly across the healthy manifold. To quantify this, we computed k-nearest-neighbour similarity in the scANVI latent space and summarized mapping specificity using weighted entropy scores (Fig. 3b,c; see Methods). Rather than forming transcriptionally isolated disease-specific clusters, the disease-associated endocrine cells remained embedded within the healthy endocrine manifold and could be interpreted relative to distinct healthy endocrine contexts resolved by HPCA (Fig. 3d). Distance calibration against held-out healthy donors further showed that the majority of the disease-associated endocrine cells remained within the range of healthy endocrine variation, with AAB-associated cells exhibiting the smallest deviations and T2D-associated cells the largest deviations from the healthy reference (Extended Data Fig. 3b).

Disease-associated beta cells showed distinct patterns of healthy-manifold occupancy. Compared with healthy beta cells, T1D-associated beta cells were enriched for immediate-early-activation, interferon-response and ER-stress neighbourhoods, whereas T2D-associated beta cells showed increased occupancy of secretory-granule and dedifferentiation-associated states. AAB-associated beta cells exhibited the strongest enrichment of INS-high and high-OxPhos neighbourhoods and reduced occupancy of INS-low states. Across diseases, INS-high represented the most frequently occupied healthy beta-cell context, consistent with multiple disease-associated beta-cell states remaining aligned to major healthy beta-cell programs (Fig. 3b). Across diseases, disease-associated alpha cells similarly localized within established HPCA alpha-cell states rather than forming disease-specific manifold regions. Relative to held-out healthy alpha cells, disease-associated alpha cells showed reduced occupancy of mature GCG-high and GCG-low programmes together with increased representation of GCG-mid, ER-stress, interferon-response and oxidative-stress neighbourhoods, with the strongest ER-stress enrichment observed in AAB-associated alpha cells (Fig. 3b). These occupancy patterns were only apparent at the level of HPCA endocrine substates, whereas conventional annotations would classify the vast majority of these cells simply as alpha or beta cells.

Weighted entropy analysis provided a complementary measure of manifold occupancy breadth relative to healthy endocrine variation. T2D-associated beta cells exhibited the highest entropy scores, indicating broader occupancy across multiple healthy beta-cell states, whereas AAB-associated beta cells showed the lowest entropy and remained localized to a more restricted region of the healthy beta-cell manifold. Alpha-cell entropy distributions were broadly similar across disease groups (Fig. 3b,c). Together, these analyses quantify disease-associated endocrine remodeling relative to healthy endocrine diversity without requiring disease-specific annotations.

These occupancy differences were accompanied by disease-context-specific transcriptional programs. T1D-associated states were characterized by increased expression of immune- and stress-associated genes, including HLA-B, B2M, IFI6, DDIT3, BIRC2 and KIAA1324, consistent with previously reported interferon-response, antigen-presentation and cellular stress programs in diabetes^69,73–76^. In contrast, T2D-associated states exhibited enrichment of genes associated with ER stress, protein folding and secretory-pathway remodeling, including HSPA5, HSP90B1, PDIA3, FKBP2, MARCH6, CASC4 and IMPAD1^69,77–82^. The AAB-associated states showed enrichment of mitochondrial, redox and proteostasis-related genes, including GLRX5, PRDX5, ATP5ME, NDUFA7, COX7C, PSMD7 and UBE2J1^83–87^. (**Fig. 3e**).

Cell–cell communication analysis (**see Methods**) identified both shared and disease-specific alterations in predicted ligand-receptor interactions across diabetes states (Fig. 3f). Canonical endocrine interactions, including INS:INSR, INS:IGF1R, GCG:GLP1R, remained detectable, whereas several pathways showed altered interaction strength including reduced SST:SSTR2 signalling in AAB and T1D. The analysis additionally identified a pan-disease loss of alpha- and beta-cell-to-macrophage CD200:CD200R1 predicted ligand-receptor interaction^88^, as well as the predicted interactions shared between T1D and T2D (PTGS2:ALOX5)^89,90^, specific to T1D (TNF:TNFRSF21)^91^, or specific to T2D (TNF:TNFRSF1A, IAPP:RAMP1)^92–94^ (Fig. 3f, Extended Data Fig. 3c).

Spatial mapping provided an independent opportunity to evaluate whether endocrine states prioritized through HPCA-based mapping could also be recovered in intact pancreatic tissue. Islets were defined using the convex hull of clusters containing at least 20 alpha cells (Extended Data Fig. 3i), and endocrine-state composition was compared between healthy and diabetic islets following spatial label transfer (Fig. 3h; see Methods). Consistent with established T1D biology, INS-expressing beta-cell states were markedly depleted within T1D islets (Extended Data Fig. 3h). In parallel, HPCA-defined interferon-response and GCG-high alpha-cell states were enriched within T1D islets, providing orthogonal support for endocrine-state assignments from dissociated-cell reference framework. Differential expression analysis of spatially resolved alpha cells further identified programs associated with inflammatory and stress-related responses, including increased expression of HLA-A, HLA-B, HLA-DRB1, GPX1 and MT2A (Fig. 3h). HPCA can therefore serve as a common coordinate system for integrating dissociated and spatial transcriptomic datasets to contextualize endocrine remodeling across diabetes-associated conditions.

Beyond contextualizing disease-associated states and spatial observations, HPCA also provides a framework for placing genetic signals into endocrine-state context. As a heuristic consistency check, we asked whether T2D-associated beta-cell programs show directional consistency with colocalized eQTL effects. We integrated published bulk islet eQTL–GWAS colocalization signals, which nominate eGenes (genes whose expression is associated with a disease-risk variant), and asked whether T2D-associated eGenes show directionally concordant expression changes across HPCA beta-cell substates (see Methods). Of 271 colocalized eGene-locus pairs queried, 208 were detectable across HPCA substates. Concordance between eQTL direction and disease-associated substate expression changes reached 81% among eGenes with compositionally stable peak substates. Both PHB (rs318095, Fig. 3j), previously identified as enriched along the T2D metabolic-stress trajectory, and FXYD2^95^ (rs529623, Fig. 3k), a T2D colocalized eGene and regulator of human beta-cell maturity, exhibited directionally concordant expression fold-changes across most annotated beta-cell substates (Extended Data Fig. 3j,k). This is consistent with their known roles in core beta-cell biology: PHB in mitochondrial integrity and oxidative metabolism^96^, FXYD2 in ion-channel-mediated regulation of insulin secretion^95^, making them tractable for follow-up across beta-cell model systems. Discordant eGenes were less likely to reflect such robust, compositionally stable associations; their mismatch may instead indicate effector mechanisms acting outside the endocrine compartment (e.g. immune or vascular islet cells) or scattered across cell types, which is a case where HPCA’s substate resolution can flag bulk eQTL signals for single-cell follow-up.

Together, these analyses illustrate how HPCA can serve as a healthy endocrine reference landscape for contextualizing diabetes-associated variation. Candidate disease-associated endocrine cells remained embedded within healthy endocrine state space, but showed disease-context-specific patterns of neighbourhood occupancy, mapping breadth and transcriptional programs, allowing remodeling-associated endocrine programs to be interpreted relative to healthy endocrine diversity.

**Extended Data Figure 3.**
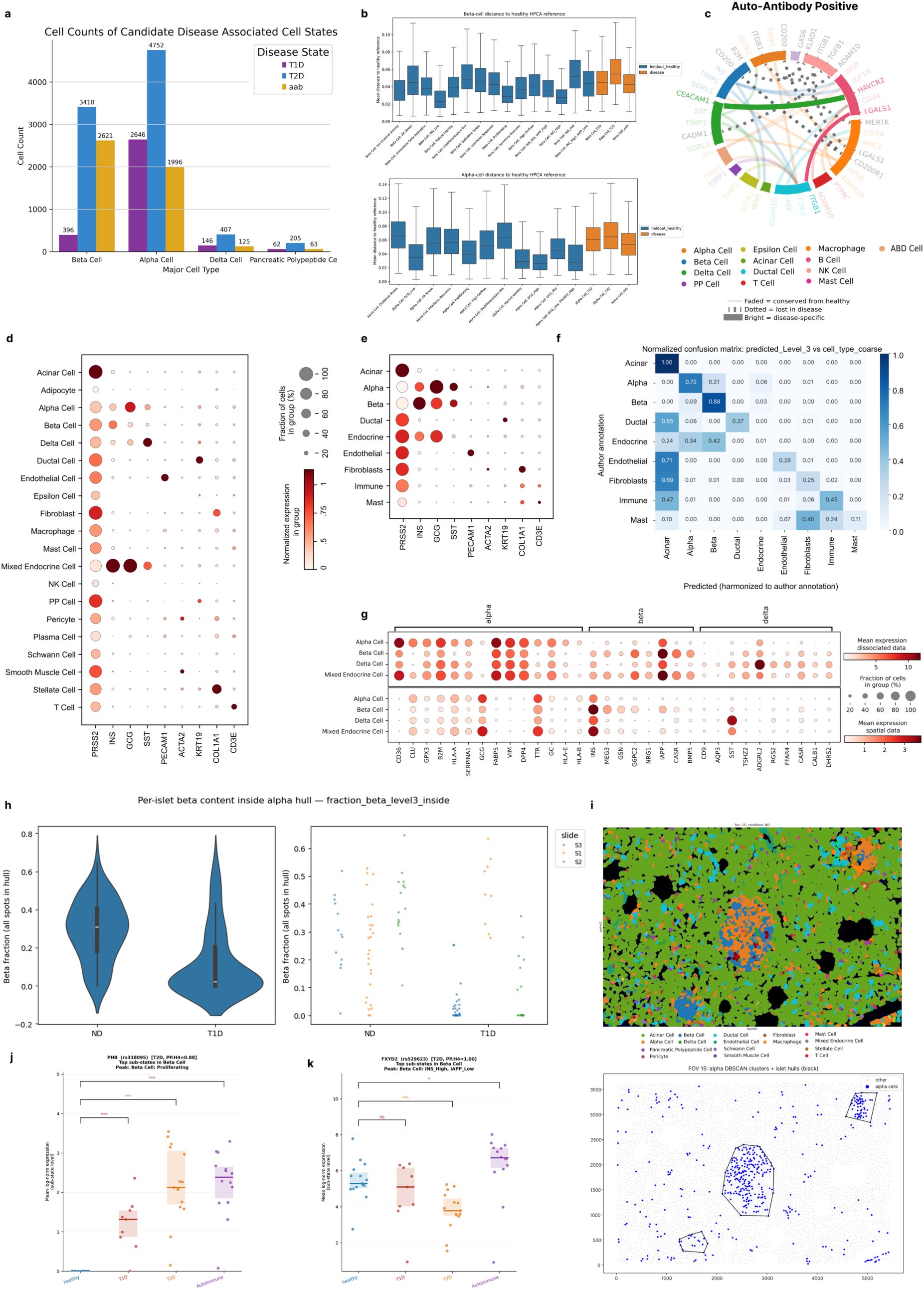
Spatial validation and characterization of diabetes-associated endocrine remodeling. **a,** Number of candidate disease-associated endocrine cells identified in T1D, T2D and autoantibody-positive (AAB) donors across major endocrine cell types. Candidate disease-associated cells were predominantly detected within alpha- and beta-cell compartments, with smaller contributions from delta and pancreatic polypeptide cells. **b,** Boxplots show distributions of cell-to-reference distances for candidate disease-associated beta-cell (top) and alpha-cell (bottom) states. Distances were calculated in the scANVI latent space relative to a healthy endocrine reference constructed from held-out healthy donors. Lower values indicate closer proximity to healthy endocrine variation. **c,** Cell–cell communication network inferred using LIANA for AAB donors. Edge width corresponds to inferred interaction strength. Faded edges indicate interactions also detected in healthy controls, dotted edges indicate interactions lost in disease and highlighted edges indicate disease-enriched interactions. **d,** Dot plot of marker-gene expression grouped by transferred Level_3 labels. **e,** Dot plot of marker-gene expression grouped by the original author annotation. **f,** Confusion matrix comparing HPCA-transferred annotations and original spatial transcriptomics annotations. Values indicate the fraction of cells assigned to each annotation class. **g,** Expression of alpha, beta and delta marker genes in dissociated and spatial data. Marker genes are defined by differential gene expression in dissociated data; markers overlapping with the spatial panel were selected. **h,** Per-islet beta-cell content within alpha-cell-defined islets in non-diabetic (ND) and T1D samples. Left, distribution of beta-cell fractions across individual islets. Right, beta-cell fractions stratified by sample. **i,** Representative spatial transcriptomics section showing HPCA-transferred cell-type annotations. Colours in upper part denote transferred cell identities. Lower part of the figure shows the alpha hulls on the same slide. **j,** Expression of PHB across healthy, T1D, T2D and autoantibody-positive beta cells. Boxplots show normalized expression values for the beta-cell substate exhibiting the highest PHB expression. Significance was assessed by two-sided Wilcoxon rank-sum tests. **k,** As in j, for FXYD2.

### Patient-level epithelial composition analysis identifies distinct injury-associated and malignancy-associated epithelial ecosystem regions

Comparing epithelial remodeling across inflammatory and malignant pancreatic diseases remains challenging because disease-associated epithelial states are often analyzed in isolation and lack a common healthy reference framework. To contextualize epithelial ecosystem organization relative to healthy pancreas, we re-integrated 555,988 HPCA exocrine epithelial cells and mapped 457,521 cells from the PDAC Atlas^97^ onto this reference using scArches^28^ (see Methods). This generated an integrated exocrine epithelial atlas of 1,013,509 cells from 28 datasets (Fig. 4a). Malignant epithelial labels from the PDAC Atlas were retained following mapping and jointly analyzed together with healthy acinar and ductal states. Donor-level epithelial profiles across 280 donors were subsequently embedded using SampleCLR^98^, followed by k-nearest-neighbour graph construction, diffusion map embedding and pseudotime-based manifold summarization (Fig. 4a,c,d; see Methods).

**Figure 4.**
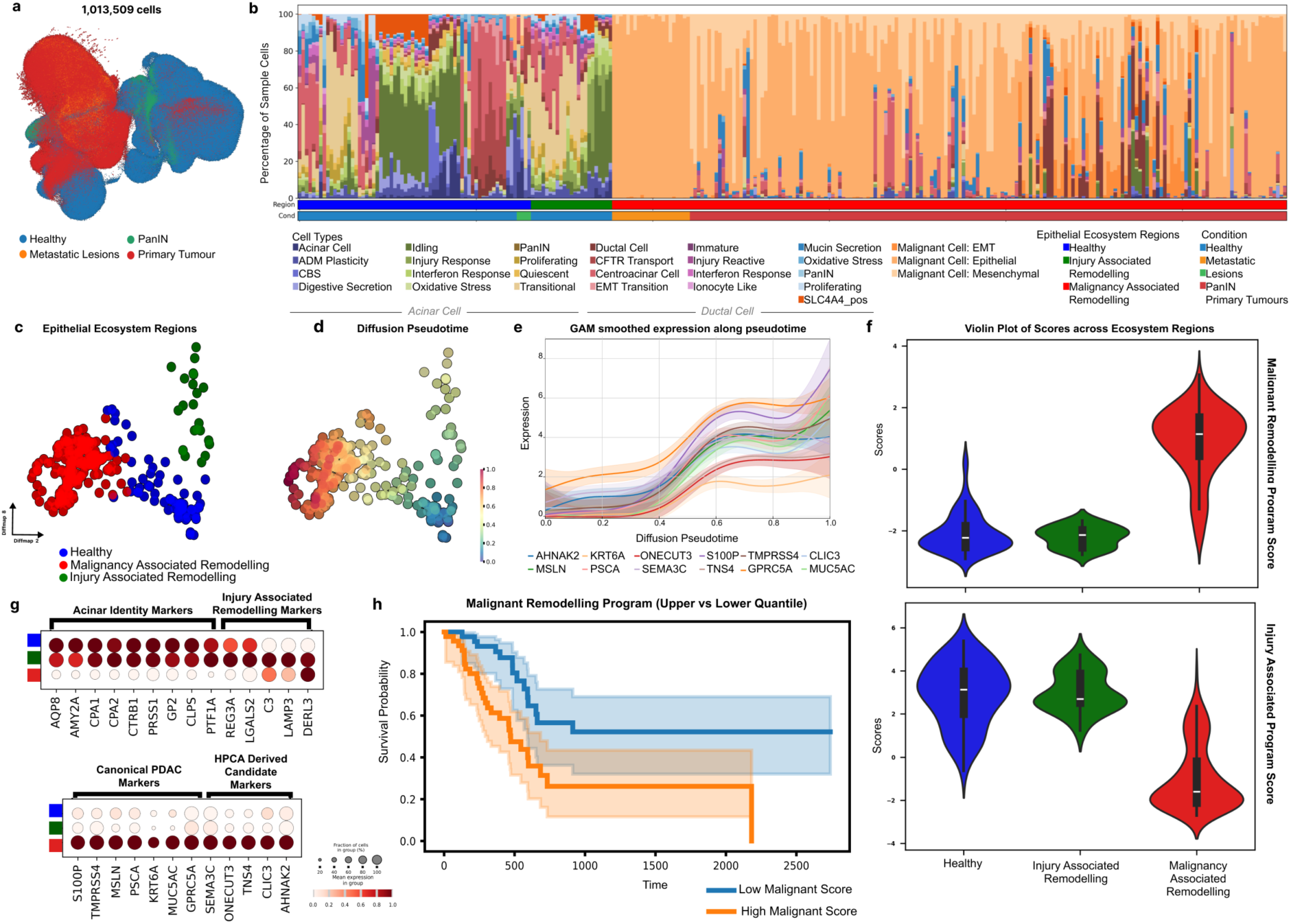
Donor-level organization of HPCA exocrine epithelial cells and mapped PDAC epithelial states reveals healthy, injury-associated and malignancy-associated remodelling regions. **a,** UMAP embedding of the integrated epithelial atlas comprising HPCA exocrine epithelial cells and PDAC Atlas epithelial cells mapped onto the healthy reference using scArches, coloured by disease condition. **b,** Donor-level epithelial composition profiles across the integrated exocrine epithelial atlas. Each bar represents one donor and shows the relative abundance of epithelial substates grouped by condition. **c,d,** Diffusion map representation of donor-level epithelial ecosystems coloured by manually annotated manifold regions corresponding to healthy homeostasis, injury associated remodeling and malignancy associated remodeling. **e,** Gene-expression dynamics along the epithelial remodeling trajectory. Generalized additive models (GAMs) were fitted to normalized expression values of selected malignancy-associated epithelial genes (AHNAK2, CLIC3, GPRC5A, KRT6A, MSLN, MUC5AC, ONECUT3, PSCA, S100P, SEMA3C, TMPRSS4 and TNS4) as a function of diffusion pseudotime. Solid lines indicate GAM-smoothed expression trends and shaded regions represent 95% confidence intervals, revealing progressive activation of malignant epithelial remodeling programs along pseudotime. **f**, Donor-level malignancy associated remodeling program scores (top) and injury associated remodeling program scores (bottom) across epithelial ecosystem regions. **h,** Expression of representative acinar identity, injury associated remodeling markers, canonical PDAC and candidate malignancy associated remodeling markers across epithelial ecosystem regions. Dot size indicates the fraction of cells expressing each gene and colour indicates mean normalized expression. **i,**Kaplan–Meier survival analysis of TCGA PAAD patients stratified by malignant-remodelling program score. Patients in the upper quartile of malignant-remodelling program enrichment exhibited poorer overall survival than those in the lower quartile. Shaded regions indicate 95% confidence intervals. Two-sided log-rank test, P = 0.00153.

Pseudotime was anchored by homeostatic acinar and differentiated ductal states to assess how disease-associated ecosystems relate to healthy pancreas organization (Fig. 4b). Diffusion pseudotime provided a one-dimensional summary of manifold organization but was not used to define ecosystem regions (see Methods). This HPCA-derived donor manifold resolved three broad compositional regions corresponding to healthy epithelial homeostasis, injury-associated epithelial remodeling, and malignancy-associated epithelial remodeling (Fig. 4c,d; see Methods). The injury-associated region was defined by donor compositions exhibiting significantly elevated injury-state scores relative to all other donor clusters (Mann–Whitney U test, P = 2.3 × 10⁻¹²). These ecosystem configurations occupied distinct topological regions of the manifold relative to the healthy reference rather than forming a single continuous compositional gradient.

Composition-level analysis further illustrated the interpretive value of the HPCA annotation framework (Fig. 4b). The injury-associated epithelial ecosystem region was enriched for acinar-cell states, including Transitional, ADM Plasticity, Injury Response and Proliferating states, together with injury-reactive ductal-cell and centroacinar-cell states. Transitional acinar states showed particularly strong enrichment alongside expansion of centroacinar and injury-reactive ductal populations. Importantly, donor profiles in this region retained substantial representation of differentiated acinar programs, including digestive secretion and quiescent acinar states, indicating preservation of mature epithelial identity alongside remodeling-associated states. The injury-associated region showed high dataset entropy (0.90), indicating contributions from multiple cohorts rather than a single-study batch effect. By contrast, the malignancy-associated epithelial ecosystem region was dominated by malignant epithelial and mesenchymal programs together with depletion of mature acinar and differentiated ductal populations. Healthy donor profiles remained enriched for homeostatic acinar compositions with minimal remodeling-associated burden.

Each epithelial ecosystem region is defined by distinct donor-level transcriptional programs, reflecting coordinated changes in epithelial composition rather than transitions of individual cell states. To characterize these programs, we generated donor-level epithelial pseudobulk profiles and identified genes whose expression varied along diffusion pseudotime within the injury-associated and malignancy-associated epithelial ecosystem regions (Fig. 4e, see Methods). Genes prioritized through correlation-based analyses and complementary differential expression analyses were subsequently summarized into composite injury-associated and malignancy-associated remodeling programs to facilitate comparison of epithelial ecosystem configurations (Fig. 4f). Donor profiles within the injury-associated epithelial ecosystem region exhibited higher injury-associated remodeling program scores and lower malignancy associated remodeling program scores than donor profiles within the malignancy-associated region (Fig. 4f).

Injury-associated donor profiles combined mature acinar secretory programs with epithelial remodeling signatures, with enrichment of digestive genes such as AQP8, AMY2A, CPA1, CPA2, CTRB1, PRSS1, GP2 and CLPS^99,100^, alongside remodeling-associated genes including REG3A, LGALS2, C3, PTF1A, LAMP3 and DERL3^101–103^ (Fig. 4h). Within the HPCA reference framework, these donor profiles were therefore characterized by concurrent representation of differentiated acinar programs and epithelial remodeling-associated signatures. In contrast, donor profiles within the malignancy-associated epithelial ecosystem region exhibited elevated malignancy associated remodeling program scores (Fig. 4f) and enrichment of genes previously associated with PDAC epithelial identity and remodeling, including S100P, TMPRSS4, MSLN, GPRC5A, MUC5AC, PSCA and KRT6A^104–110^ (Fig. 4g). Recovery of these established PDAC-associated genes provides an internal positive control that the HPCA framework captures known disease-associated epithelial programs. Additional malignant region associated genes included SEMA3C, TNS4, CLIC3, AHNAK2, and ONECUT3, previously implicated in epithelial plasticity and tumor remodeling across multiple cancer types^88,111–116^. While several of these genes have been reported individually in prior studies, their coordinated enrichment within the malignancy-associated epithelial ecosystem region suggests a distinct remodeling-associated epithelial program and identifies candidate regulators for future investigation (Fig. 4g).

To determine whether epithelial remodeling programs identified through HPCA were observable beyond the discovery datasets, we evaluated their expression in TCGA PDAC transcriptomes. Tumors with higher expression of the HPCA-derived malignancy associated remodeling program exhibited poorer overall survival than tumors with lower program expression (upper-versus-lower quartile comparison; two-sided log-rank test, P = 0.0015; Fig. 4h; see Methods). Consistent with this observation, malignancy associated remodeling scores were associated with worse survival in Cox proportional hazards models (hazard ratio, 1.52; 95% confidence interval, 1.01–2.29; P = 0.043). Several candidate remodeling-associated genes prioritized through the HPCA framework, including AHNAK2, TNS4, SEMA3C and CLIC3, also demonstrated significant survival stratification in TCGA PDAC cohorts (Extended Data Fig. 4a-d). Although these analyses do not establish prognostic utility, they provide orthogonal support that epithelial ecosystem programs identified through HPCA are reproducibly detectable in an independent cohort.

Collectively, these analyses reveal that pancreatic epithelial remodeling is organized into distinct ecosystem configurations relative to healthy epithelial homeostasis. Injury-associated ecosystems retain substantial differentiated acinar identity while acquiring remodeling-associated states, whereas malignancy-associated ecosystems exhibit coordinated loss of homeostatic epithelial programs together with expansion of malignant epithelial and mesenchymal populations. By providing a common healthy epithelial coordinate system, HPCA enables direct comparison of these ecosystem architectures across independent disease cohorts without assuming a linear progression model.

**Extended Data Figure 4.**
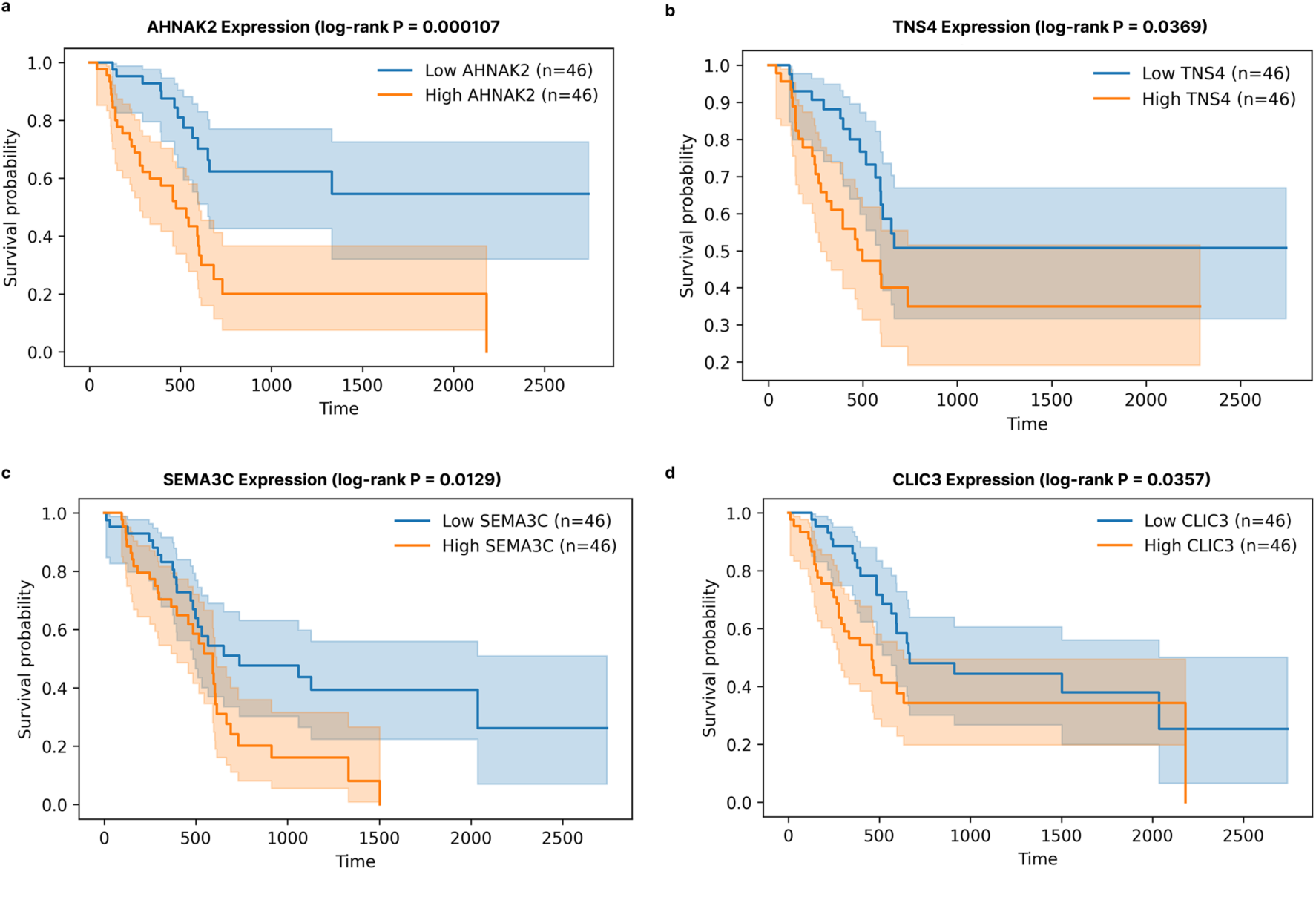
Survival associations of candidate malignant remodeling genes in TCGA PDAC. **a–d,** Kaplan–Meier survival analyses of TCGA PDAC patients stratified by expression of candidate malignant remodeling genes identified from the HPCA–PDAC epithelial ecosystem analysis. Patients were divided into upper- and lower-expression groups (n = 46 patients per group). a, AHNAK2. b, TNS4. c, SEMA3C. d, CLIC3. Shaded regions indicate 95% confidence intervals. P values were calculated using two-sided log-rank tests.

### Cross-species endocrine mapping reveals convergent stress-remodeling programs across murine diabetes models

Experimental mouse models are central to diabetes research, but the extent to which individual models represent human endocrine remodeling programs remains difficult to quantify. HPCA enables cross-species comparison of endocrine state representation using 193,453 endocrine cells from the Hrovatin et al. mouse islet atlas (MIA)^117^, spanning nine scRNA-seq datasets, 56 samples and multiple diabetic perturbation models, including autoimmune (NOD), metabolic (db/db) and streptozotocin-induced beta-cell ablation systems. Mouse endocrine cells were projected onto the HPCA endocrine manifold using a weighted k-nearest-neighbour (wkNN) mapping framework (Fig. 5a,b; see Methods). To quantify representation of HPCA endocrine states across models, we computed neighbourhood-based presence scores^118^ that measured the extent to which mapped mouse endocrine cells occupied HPCA endocrine neighbourhoods (see Methods). Presence scores therefore quantify representation of HPCA transcriptional neighbourhoods by mapped mouse cells.

**Figure 5.**
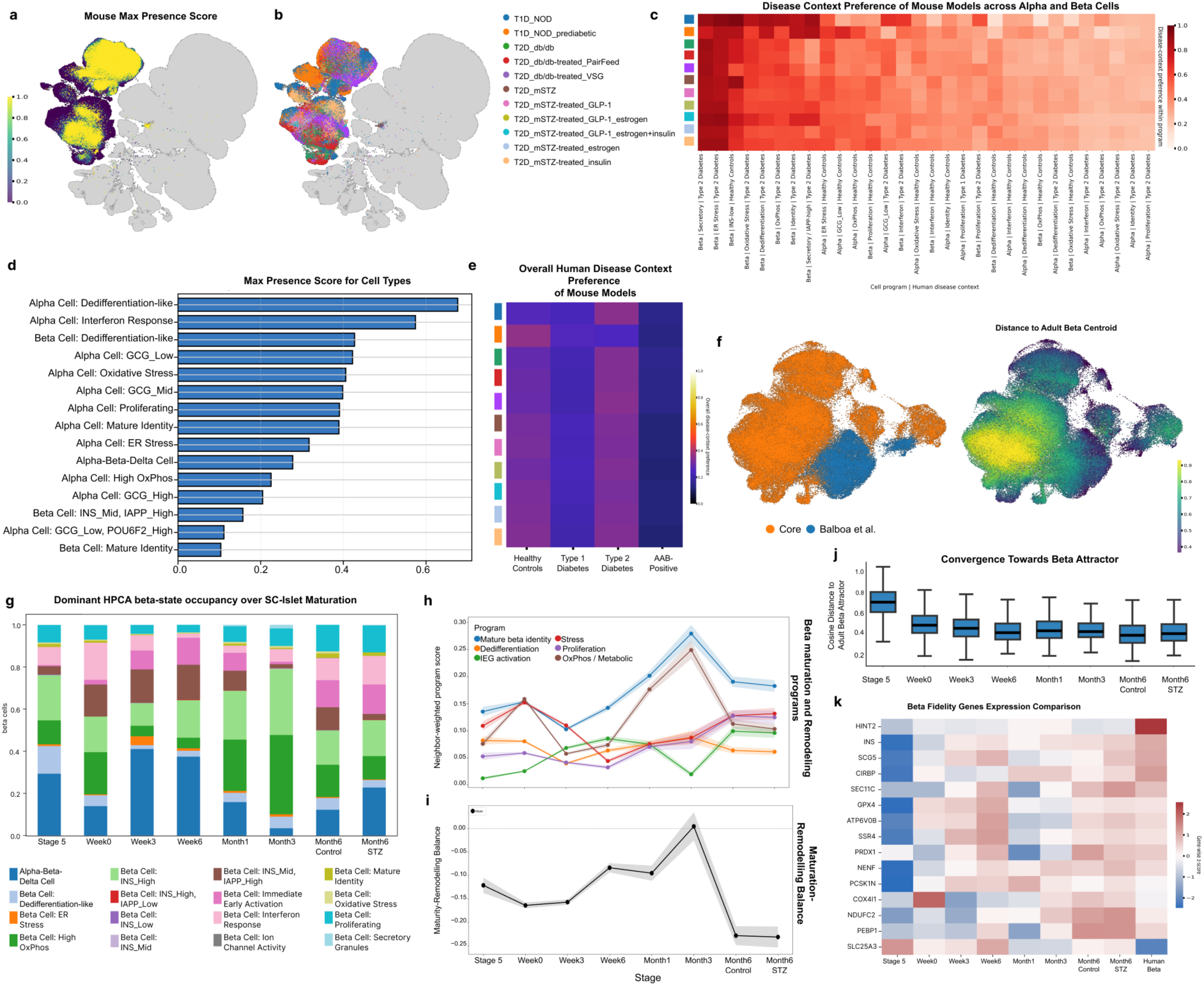
Endocrine benchmarking reveals conserved stress-remodelling programs across murine diabetes models and incomplete adult human beta-cell identity in stem-cell-derived islets. **a,** Maximum mouse-model presence score, indicating the highest presence score assigned to each human endocrine neighbourhood across all mapped mouse-model conditions. **b,** Maximally represented mouse-model condition, indicating which mouse model contributed the highest presence score for each human endocrine neighbourhood. **c,** Disease-context preference scores of murine diabetes models across conserved human alpha- and beta-cell programs. Columns represent HPCA endocrine states stratified by human disease context, including healthy controls, T1D, T2D and autoantibody-positive (AAB) donors. **d,** HPCA endocrine states most consistently represented across murine diabetes models, ranked by mean presence score across mapped conditions. Stress-adaptive endocrine programs, including interferon-response, oxidative-stress, ER-stress and dedifferentiation-associated states, showed broader representation than mature endocrine identity states. **e,** Overall human disease-context preference across murine diabetes models, summarizing relative similarity to healthy, T1D, T2D and AAB endocrine programs. For each mouse model, mean HPCA neighbourhood presence scores were aggregated across all represented alpha- and beta-cell programs for each human disease context (healthy control, T1D, T2D and AAB) and normalized to generate relative context-preference scores. **f,** Projection of SC-islet beta cells onto the HPCA beta-cell manifold. Right, HPCA beta-cell reference coloured by cosine distance to the adult beta-cell attractor centroid. Left, mapped SC-islet beta cells coloured by dataset of origin. Dominant HPCA beta-cell neighbourhood occupancy across SC-islet maturation stages. **g,** Stacked bar plots show the fraction of SC-islet beta cells assigned to each HPCA beta-cell state based on dominant neighbourhood mapping. **h,** Stage-wise trajectories of conserved beta-cell programs in SC-islets. Neighbor-weighted program scores were calculated by projecting SC-islet beta cells onto HPCA beta-cell reference states and aggregating related reference programs, including mature beta-cell identity, dedifferentiation, immediate-early gene activation, stress-associated, proliferative and oxidative phosphorylation-associated states. Lines indicate stage-wise means and shaded regions indicate ±SEM across mapped SC-islet beta cells. **i,** Maturity-remodeling balance across SC-islet differentiation stages, calculated as the mature beta-cell identity score minus the combined remodeling/plasticity score (dedifferentiation, immediate-early activation, stress and proliferation programs). Higher values indicate greater enrichment of mature beta-cell identity relative to remodeling-associated states. Lines indicate stage-wise means and shaded regions indicate ±SEM across mapped SC-islet beta cells. **j,** Distribution of cosine distances between SC-islet beta cells and the adult HPCA beta-cell attractor across differentiation stages. Lower values indicate greater similarity to the adult beta-cell reference state. Boxplots summarize the median and interquartile range, with whiskers extending to 1.5× the interquartile range. **k,** Expression of representative adult beta-cell fidelity genes across SC-islet maturation stages and adult HPCA beta cells. Heatmap values represent scaled gene expression.

Across mouse models, representation was concentrated within a subset of HPCA endocrine neighbourhoods rather than uniformly distributed across the healthy endocrine reference landscape (Fig. 5d). The highest-scoring HPCA neighbourhoods were dominated by alpha-cell programs, including dedifferentiation-like, interferon-response, GCG-low, GCG-mid, oxidative-stress and proliferative states, together with beta-cell dedifferentiation-like states. In contrast, several beta-cell neighbourhoods, including Mature Identity and INS_Mid, IAPP_High, showed comparatively lower representation (Fig. 5d). Alpha-cell-associated neighbourhoods occupied the majority of the fifteen most highly represented HPCA endocrine states, indicating that diverse alpha-cell transcriptional contexts are broadly recovered across murine diabetes models. Multiple independent mouse systems mapped to overlapping alpha-cell neighbourhoods annotated as interferon-response, oxidative-stress, proliferative, ER-stress and dedifferentiation-associated states. Many of these neighbourhoods corresponded to previously described endocrine stress-response and remodeling-associated programs^15,117^, providing an internal positive control for the cross-species mapping framework. These results suggest that commonly used diabetes models preferentially capture endocrine stress- and remodeling-associated transcriptional programs, whereas mature endocrine identity states are represented less consistently across models.

We next stratified endocrine neighbourhoods according to their relative enrichment in healthy controls, T1D, T2D and autoantibody-positive (AAB) donors within the human disease-mapping analyses described above, and quantified disease-context preference within each represented neighbourhood (Fig. 5c). Rather than assigning mouse cells directly to disease labels, this approach places model-derived endocrine cells within HPCA neighbourhoods that exhibit varying degrees of association with different human disease contexts. Program-level context preferences differed substantially across endocrine neighbourhoods (Fig. 5c). T2D-associated beta-cell programs, including Secretory, ER-Stress, Oxidative-Stress and Dedifferentiation neighbourhoods, were among the most consistently represented contexts across models, whereas several alpha-cell neighbourhoods showed stronger association with healthy-control contexts. Despite these program-level differences, aggregation across all represented alpha- and beta-cell programs revealed a broadly similar overall pattern across mouse models (Fig. 5e). Across models, overall context-preference summaries showed partial alignment with healthy-control and T2D-associated endocrine contexts, while AAB-associated contexts contributed less across all mouse systems (Fig. 5e). Importantly, neither autoimmune nor metabolic mouse models exhibited exclusive alignment with their corresponding human disease contexts. For example, NOD^119^, models retained substantial representation across healthy-control and T2D-associated contexts despite their autoimmune disease background, indicating that HPCA context-preference scores capture representation of shared endocrine neighbourhoods rather than direct disease equivalence, even when overall program distributions exhibit relative similarity to human T1D profiles.

Because human disease contexts differ in their distribution across HPCA endocrine programs, we additionally compared mouse-model program-occupancy distributions with the corresponding distributions observed across human disease contexts (Extended Data Fig. 5a). Prediabetic NOD samples showed the strongest similarity to the human T1D program distribution (cosine similarity = 0.50), whereas db/db and mSTZ-derived models exhibited comparable similarity to healthy-control, T1D and T2D distributions. No mouse model showed exclusive similarity to a single human disease context, and similarity scores remained broadly distributed across contexts. More broadly, similarity scores were distributed across multiple human contexts rather than concentrating within a single disease category. These findings suggest that individual mouse models capture overlapping subsets of endocrine remodeling programs that are shared across human disease contexts, rather than uniquely representing a single human disease state.

Together, these analyses demonstrate that experimental diabetes models capture overlapping but incomplete subsets of the human endocrine landscape. Rather than identifying a single model that best recapitulates a given disease, HPCA enables quantitative benchmarking of which endocrine neighbourhoods and remodeling-associated programs are represented across model systems. This framework highlights both shared endocrine stress programs that are reproducibly recovered across models and regions of the human endocrine landscape that remain comparatively underrepresented, thereby informing model selection for specific biological questions and identifying areas where complementary experimental systems may be required.

### The HPCA defines an adult beta-cell reference landscape for benchmarking stem-cell-derived islets

As with mouse models, assessing the fidelity of stem-cell-derived beta cells remains challenging owing to the lack of comprehensive adult human reference landscapes. To demonstrate the broader utility of HPCA for benchmarking model systems against adult human cell states, we mapped 43,790 quality-controlled stem-cell-derived islet (SC-islet) cells from Balboa et al.^41^ onto the HPCA reference, spanning differentiation stages from in vitro Stage 5 through week 6, together with engrafted islets collected 1, 3 and 6 months post-transplantation, and restricted downstream analyses to query cells assigned to beta-cell-associated HPCA neighbourhoods. The scale and diversity of HPCA healthy human donors enabled construction of a comprehensive adult beta-cell reference landscape and an adult beta-cell centroid representing a consensus transcriptional context across healthy individuals (see Methods).

Dominant-neighbourhood analysis showed that SC-islet beta cells occupied a heterogeneous mixture of HPCA beta-cell neighbourhoods across differentiation stages, including mature beta identity, high-OxPhos, INS-high, INS-mid/IAPP-high, dedifferentiation-like, immediate-early activation, stress-associated and proliferative states (Fig. 5g). Rather than mapping predominantly to a single adult beta-cell context, SC-islet beta cells distributed across multiple reference neighbourhoods, indicating representation of diverse adult beta-cell transcriptional contexts within the SC-islet system.

To quantify these neighbourhood assignments, we summarized query-to-reference connectivity into maturation- and remodeling-associated HPCA programs. Representation of mature beta-cell programs increased across differentiation and was most strongly enriched at Month3, accompanied by increased occupancy of high-OxPhos neighbourhoods (Fig. 5h). The balance between predefined mature and remodeling-associated HPCA state groups also varied across stages, with Month3 exhibiting the strongest relative representation of mature reference programs. Later stages showed increased representation of proliferative, stress-associated and immediate-early activation neighbourhoods (Fig. 5i,j). These observations provide a quantitative summary of how adult beta-cell reference programs are represented across SC-islet stages, without requiring assumptions regarding developmental trajectories or maturation pathways.

We next asked whether neighbourhood-level representation corresponded to global similarity to the adult beta-cell reference landscape (Fig. 5f). Using the HPCA-derived adult beta-cell centroid, we calculated the mean transcriptomic distance of SC-islet beta cells to the adult reference across differentiation stages(Fig. 5j). Similarity to the adult centroid increased from Stage5 through Week6 and remained relatively high at subsequent stages, with Month6_CTRL exhibiting the closest overall correspondence to the adult reference centroid (Fig. 5j). Notably, stages exhibiting strong global similarity to the adult reference landscape continued to display heterogeneous occupancy across multiple HPCA neighbourhoods, indicating that centroid-level similarity and neighbourhood-level representation capture complementary aspects of reference-state correspondence.

Finally, we examined transcriptional programs associated with proximity to the adult beta-cell reference centroid. SC-islet beta cells increasingly represented components of the adult beta-cell fidelity program during differentiation, including INS and genes associated with secretory function, protein handling, redox homeostasis and mitochondrial metabolism. Several centroid-associated genes, including HINT2, SCG5, CIRBP, SEC11C, GPX4, SSR4, PRDX1, PCSK1N and SLC25A3, were consistently lower or more variably expressed in SC-islet beta cells relative to adult reference beta cells (Fig. 5k). These genes define components of the adult beta-cell reference program that remain incompletely represented in the SC-islet system and provide candidate targets for optimizing SC-islet differentiation and maturation protocols.

Together, these analyses position SC-islet beta cells within a donor-derived adult beta-cell reference landscape and demonstrate how HPCA can be used to compare neighbourhood occupancy, reference-program representation and global transcriptional similarity across model systems within a common coordinate system.

**Extended Data Fig. 5.**
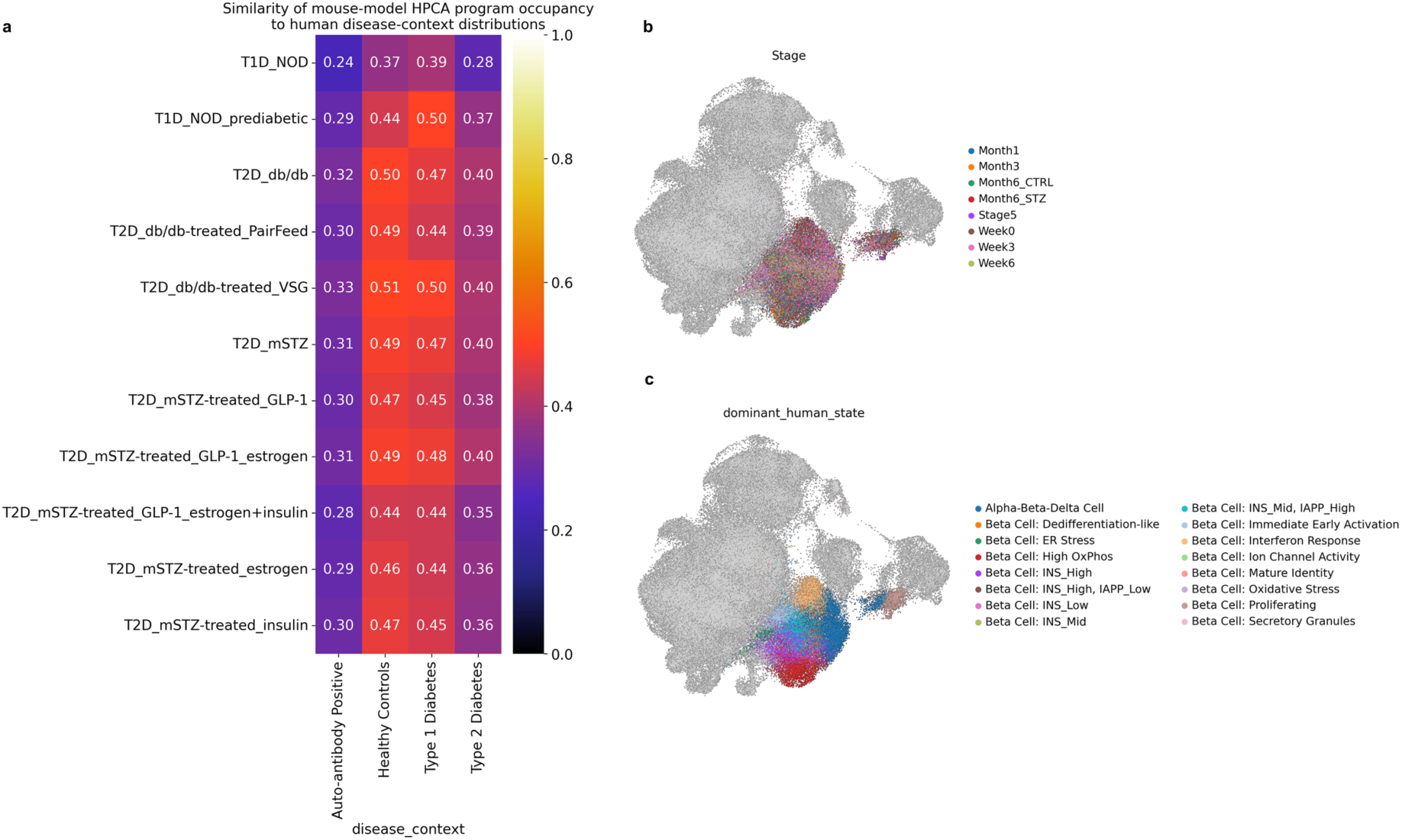
Cross-species benchmarking of murine diabetes models and stem-cell-derived islets within the HPCA endocrine reference landscape. **a,** Similarity between endocrine program occupancy distributions observed across murine diabetes models and human disease contexts. Mouse endocrine cells from the mouse islet atlas were mapped onto the HPCA endocrine manifold, and occupancy distributions across HPCA endocrine neighbourhoods were compared with those observed in human healthy-control, autoantibody-positive (AAB), type 1 diabetes (T1D) and type 2 diabetes (T2D) donors. Values indicate cosine similarity between mouse-model and human disease-context program distributions. **b,** Projection of stem-cell-derived islet (SC-islet) beta cells from Balboa et al. onto the HPCA beta-cell reference manifold. Cells are coloured according to differentiation stage, including Stage 5, Week 0, Week 3, Week 6, Month 1, Month 3, Month 6 control (CTRL) and Month 6 streptozotocin-treated (STZ) samples. Grey points indicate HPCA reference beta cells. **c,** Dominant HPCA beta-cell neighbourhood assigned to mapped SC-islet beta cells. Colours indicate the dominant human reference neighbourhood inferred from weighted nearest-neighbour mapping. SC-islet beta cells occupy multiple adult beta-cell reference states, including mature identity, INS-high, INS-mid/IAPP-high, oxidative-stress, ER-stress, dedifferentiation-like, proliferative and immediate-early activation-associated neighbourhoods.

## Discussion

We have built the HPCA, a high-resolution healthy Human Cell Atlas reference framework for interpreting cellular diversity, disease remodeling, and experimental model systems across the human pancreas. By integrating 815,126 single-cell and single-nucleus transcriptomes from 109 donors across diverse studies, technologies and tissue-processing protocols, HPCA captures broad healthy pancreatic variation while resolving 94 cell types and transcriptional states across endocrine, exocrine, immune, endothelial and stromal compartments. This granularity enables the atlas to move beyond broad cell-type annotation and instead define a healthy transcriptional coordinate system against which new datasets can be mapped, compared and interpreted. Beyond serving as a reference for label transfer, HPCA establishes consensus cellular identities and marker programs across studies, providing a common vocabulary for interpreting pancreatic cellular variation. Similar efforts in other tissues have demonstrated that integrated reference atlases can resolve cellular heterogeneity, define robust marker genes and establish common frameworks for biological interpretation across studies and experimental systems^11,12^.

A central impact of HPCA is that it enables pancreatic disease states to be interpreted relative to the structure of healthy cellular diversity rather than only within disease datasets themselves. In diabetes, candidate disease-associated endocrine cells remained embedded within the healthy endocrine manifold, but showed condition-specific occupancy of stress, interferon-response, secretory-remodeling, oxidative-stress and dedifferentiation-associated neighbourhoods. Similarly, epithelial remodeling across healthy and PDAC samples organized at the donor level into distinct homeostatic, injury-associated, and malignancy-associated ecosystem regions. These analyses illustrate how healthy reference atlases can serve as coordinate systems for constructing disease manifolds, enabling disease-associated remodeling to be quantified relative to healthy cellular programs rather than inferred from disease datasets in isolation. Such frameworks enable quantitative comparison of remodeling across cohorts, diseases and experimental systems. Thus, these findings support a view of pancreatic disease as remodeling of pre-existing healthy cellular programs, with disease-associated states occupying distinct regions of a broader healthy-to-disease transcriptional landscape rather than forming entirely separate cellular identities^24–26^.

The depth of HPCA additionally enabled identification and contextualization of rare endocrine populations, including a polyhormonal αβδ state. Rare polyhormonal endocrine populations have previously been reported during pancreatic development, regeneration and disease-associated contexts^41–43^. Although mixed-lineage endocrine populations remain challenging to distinguish from technical artifacts^120^, the scale of HPCA enabled detection and prioritization of rare cellular states that may be missed in individual studies^11^. Importantly, HPCA is constructed from histologically non-diseased donor tissue, which may not always correspond to the absence of subclinical pathology. Thus, rare states such as polyhormonal αβδ cells could reflect undetected physiological or pathological remodeling in a subset of donors. Nevertheless, their detection across multiple datasets and individuals suggests that they are present in a meaningful proportion of the broader human population, highlighting the value of integrated references for prioritizing rare candidate cellular states for future investigation.

HPCA also provides a translational benchmarking platform for evaluating experimental systems against adult human reference states. Cross-species mapping showed that murine diabetes models preferentially represented conserved endocrine stress-remodeling programs rather than uniformly recapitulating human disease-associated endocrine states. Prior studies have highlighted important biological differences between mouse and human endocrine systems and diabetes progression^20^. Mapping stem-cell-derived islets onto the HPCA beta-cell landscape similarly revealed partial acquisition of adult beta-cell identity alongside persistent representation of remodeling, stress and proliferative programs. Previous studies have likewise shown that stem-cell-derived beta cells frequently retain immature features despite expression of canonical endocrine markers^121,122^. Together, these analyses demonstrate how reference atlases can move beyond annotation to provide quantitative benchmarks for translational relevance, enabling systematic assessment of which human cellular programs are faithfully recapitulated by experimental systems and which remain incompletely modeled.

Several limitations should be considered. First, although benchmarking supported scANVI as the optimal integration strategy, integration-based references remain sensitive to dataset composition, suspension type, batch covariates and annotation structure^27,28^. Consequently, rare populations, transcriptional continua and neighbourhood boundaries may be refined as additional cohorts, technologies and annotations are incorporated. HPCA should therefore be viewed as an evolving rather than definitive representation of healthy pancreatic diversity. Moreover, while HPCA captures substantial donor, demographic and technical diversity, fully representing healthy pancreatic variation across age, ancestry and environmental exposures remains an ongoing challenge that future atlas iterations, including efforts through the HCA Ancestry Network, will help to address. Second, disease-associated cells identified through mapping uncertainty and neighbourhood occupancy should be interpreted as candidate remodeling states rather than definitive disease-specific cell types. Cells can occupy disease-enriched regions of the healthy manifold without representing stable disease-specific identities, and HPCA therefore prioritizes candidate remodeling states for further investigation rather than defining new disease-specific cell types. Third, donor-level epithelial manifolds and pseudotime analyses summarize ecosystem organization relative to healthy reference states but do not establish lineage relationships, temporal progression or causal transition^123^. Longitudinal sampling and matched precursor-to-tumour datasets will be required to determine how these ecosystem configurations relate to disease evolution. Fourth, inferred ligand–receptor interactions are hypothesis-generating and require spatial, proteomic or functional validation. Consequently, these interactions should be viewed as candidate signaling relationships rather than direct evidence of active cell–cell communication. Finally, spatial, genetic and TCGA-based analyses provided orthogonal support for HPCA-derived observations but remain limited by targeted spatial panels, bulk-tissue resolution and the absence of paired single-cell genotype or longitudinal data. Future multimodal datasets integrating transcriptomic, spatial, genetic and functional measurements will be required to move from reference-based prioritization to mechanism.

Looking forward, HPCA provides a foundation for increasingly multimodal and disease-aware pancreas atlases integrating spatial context, genetic variation, clinical phenotypes and longitudinal disease sampling. More broadly, this study illustrates how healthy reference atlases can function as evolving transcriptional coordinate systems that connect normal tissue variation, disease remodeling and experimental perturbation within a unified analytical framework. As such resources mature across tissues, they may provide a general strategy for interpreting cellular states, transitions and therapeutic responses across human biology^10–12^.

## Methods

### Collection and Preprocessing of Human Datasets

We collected 12 public and pre-published available single-cell and single-nucleus RNA sequencing (scRNA-seq and snRNA-seq) datasets^6,8,9,15–21^ from GEO and Consortiums, ensuring the inclusion of metadata from associated manuscripts (Supplementary Table 1). For preprocessing, we applied consistent quality control across all datasets. To remove potential outliers, cells with total counts below the 5th percentile or above the 95th percentile were excluded. Genes expressed in less than 10 cells were removed, as well as cells expressing fewer than 400 genes. This stringent preprocessing ensured high-quality data for downstream analysis. Furthermore, we collected 5 additional publicly available studies^68–72^ for the extension of the core atlas. The preprocessing was done in the similar fashion to the core atlas.

### Metadata collection and harmonization

Metadata was collected in accordance with the Human Cell Atlas Tier 1 metadata requirements^124^.

### Read alignment and count matrix generation

To ensure consistency across datasets, all samples were harmonized to the human reference genome (GRCh38). For most datasets, publicly available gene expression count matrices already aligned to GRCh38 were used directly. However, three datasets—GSE84133, GSE101207, and GSE114297—were reprocessed from raw sequencing data. These datasets were either originally aligned using non-standard pipelines, provided only as FASTQ files, or aligned to earlier genome builds. In these cases, reads were realigned to GRCh38 to generate standardized gene expression count matrices for downstream analysis.

### Data Normalization

Raw count matrices from each dataset were normalized using Scanpy’s normalize_total function (target sum = 1 × 10⁶) followed by logarithmic transformation (log1p). To reduce the impact of extreme expression values and improve comparability across datasets.

### Principal Component Regression and Theil’s U for Selection of Batch Covariate

To identify the primary source of technical variation, we evaluated candidate covariates using principal component regression (PCR) and Theil’s uncertainty coefficient (Theil’s U). Principal components were computed on log-normalized expression data, and the variance explained (R²) by each covariate was quantified across the top components using linear models (PC ∼ covariate). In parallel, dependence between covariates and transcriptional structure was assessed using Theil’s U, computed on discretized PC scores. The covariate showing the strongest and most consistent association across both metrics was selected as the batch variable for downstream integration. For datasets exhibiting nested batch effects, we made an informed decision to further stratify data at the donor level.

### Annotation Harmonization

Cell type annotations were harmonized across datasets to establish a consistent reference. Eight datasets with existing author annotations were standardized to a unified ontology, while the remaining datasets were manually annotated and aligned to the same schema. Harmonized Level 2 annotations were used for downstream integration and benchmarking.

### Feature Selection for Integration

Highly variable genes (HVGs) were selected by jointly considering scRNA-seq and snRNA-seq datasets. HVGs were computed separately for each modality, and both the union and intersection were evaluated at multiple thresholds (1,000, 2,000, 3,000, 4,000, and 8,000 genes). Gene sets were further augmented with curated marker genes to preserve biological signals. Based on preliminary integration and benchmarking, the intersection of 3,000 HVGs supplemented with marker genes provided the optimal balance between batch correction and biological conservation for the preliminary integration.

### Integration benchmarking

Integration performance was evaluated across multiple methods, including scVI, scANVI, DRVI, Harmony, sysVI, and scPoli as well as different count preprocessing strategies (raw counts, binned gene expression^125^) Methods were benchmarked using standardized metrics from scIB, assessing both batch mixing and biological conservation. The selected feature set (3,000 HVGs plus marker genes) consistently yielded the best overall performance.

### Community-driven annotation via HCA Cell Annotation Platform

To obtain high-confidence cell type annotations, the integrated dataset was uploaded to the HCA Cell Annotation Platform (CAP). Annotators were provided with three different resolutions of leiden clustering. Independent annotations were provided by four expert laboratories within the Bionetwork. These annotations were subsequently harmonized across contributors through consensus-based curation, resulting in the final Level 3 cell type annotations. From these Level 3 cell type labels, a coarser label hierarchy was created encompassing broader lineage definitions.

### Cell State Annotations

Cell states were defined within major pancreatic lineages, including exocrine (acinar and ductal) and endocrine (alpha, beta, delta and pancreatic polypeptide cells) compartments. Gene program–based scoring was performed to capture functional states, followed by within-dataset and suspension-type z-normalization. Cells were assigned to the state with the maximum score if the corresponding z-score exceeded 1.5 standard deviations; otherwise, cells retained their Level-3 identity. This approach resulted in 94 final labels at Level 4, representing a mixture of refined cell types and transcriptional states at the highest annotation resolution.

### Final Integration

To generate the final integrated representation, we constructed a feature set comprising 500 global highly variable genes and 100 lineage-specific HVGs per Level 3 cell type. The dataset was then re-integrated using scANVI with Level 4 annotations as supervision to obtain the final embedding (scArches v0.6.1)^28^.

### Distance between Cell states in latent space

To analyse distances between cell states, we computed euclidean distances between centroids of cell states (Level_4 labels) in the latent embedding space before and after reintegration.

### Level 4 Entropy Score

To quantify heterogeneity and potential batch-driven variability across cell populations, we computed Shannon entropy for each Level 4 cell type with respect to donor, dataset, and suspension type. Entropy values were calculated based on the distribution of cells across these covariates and averaged to obtain a final entropy score per cell type.

### Ambient RNA assessment

To quantify ambient RNA contamination, we defined lineage-specific ambient RNA gene sets derived from highly expressed genes in major source populations. Cells were scored for ambient RNA signatures using log-normalized expression values. For each cell type not corresponding to the source of the ambient signal, cells within the top 10% of scores were classified as exhibiting elevated ambient RNA. Ambient RNA contamination was evaluated across key sources, including acinar, alpha, and beta signatures. In total, 165,573 cells displayed elevated ambient RNA signals. These cells were retained in the atlas and flagged to preserve data completeness, while downstream analyses most sensitive to cross-lineage marker expression, including differential gene expression analyses involving endocrine populations, were interpreted with this annotation and repeated where appropriate after excluding flagged cells.

### Final quality control

To restrict the final atlas to high quality cells, a post hoc quality control encompassing lineage informed cell filtering based on 4 median absolute deviations of number of genes and number of counts, as well as doublet filtering based on doublet detection by doubletdetection^126^ (v. 4.3) was performed. Cells not passing doublet detection in the alpha-beta-delta cell cluster were retained if they passed all other quality control methods due to the stringent, additional quality control performed on the cell population. These quality control filtering steps removed 59,807 leaving 815,126 high quality cells.

### Metadata covariate variance contribution analysis

Metadata variance contribution analysis was performed as previously described in Sikkema et al.^11^.

### Differential gene expression analysis (Obesity)

Cells per Level 3 cell type were pseudobulked and differential gene expression was computed on BMI bins based on WHO weight classification^127^ between healthy and obese bins with PyDEseq2^128^. Dataset and BMI_category were considered in the differential gene expression formula. Overrepresentation analysis was performed using PYGSEA, using all expressed genes in the cell type as background.

### Cell cell communication analysis

Cell-cell communication (CCC) was inferred using LIANA^51^ v1.7.1. LIANA was run independently on each dataset of the core healthy pancreas atlas rather than on the pooled atlas, to avoid confounding interaction scores with dataset-level batch effects in gene expression. Interactions were analysed at Level 3 cell-type resolution, a minimum of 10 cells per cell type was required per dataset. Because immune cells are underrepresented within individual datasets, immune cells were pooled at the condition level (T1D, T2D, AAB, healthy) across all datasets prior to analysis, using Level 3 annotations. Each per-dataset LIANA run combines that dataset’s endocrine and exocrine cells with the full condition-level immune pool, such that variation in immune interaction scores across datasets is driven by the endocrine and exocrine side only. Only interactions involving at least one immune cell type were retained from these runs. Interaction strength was quantified using the magnitude_rank score produced by LIANA’s rank_aggregate method, lower values indicate stronger interactions. To identify interactions enriched in disease, we computed for each ligand-receptor pair the difference in mean magnitude_rank between disease samples and matched healthy samples; pairs with a negative difference (disease − healthy < 0) were considered disease-enriched. Novel disease-specific interactions, defined as those consistently active in disease but not detected at the same level in healthy tissue.

### Spatial data analysis

Labels of the healthy atlas were mapped to spatial data published in Melton, Jimenez et al.^9^ using TACCO^40^. Labels were transferred in two rounds: on Level 3 and Level 4 labels and verified visually on a per slide level and compared against coarse author annotations. Islet spots in the data were identified using DBSCAN^129^, as implemented in scikit learn, spots with at least 20 α-cells were considered islets. An islet mask was calculated based on the convex hull of the alpha cells in each islet. Compositional differences within islets were tested with Mann-Whitney U, P-Values were multiple testing corrected with Benjamini-Hochberg.

### Quality-control analyses of αβδ cells

To evaluate whether αβδ cells could be explained by ambient RNA contamination or technical artifacts, distributions of alpha-associated ambient scores, beta-associated ambient scores, ribosomal transcript fractions, and mitochondrial transcript fractions were compared between αβδ cells and reference endocrine populations. Global distributional differences were quantified using the two-sample Kolmogorov–Smirnov (KS) statistic. For donor-level inference, median values for each metric were calculated separately within each donor and cell population. Paired differences between αβδ cells and the corresponding reference population were assessed across donors using two-sided Wilcoxon signed-rank tests. Alpha-associated ambient scores were compared against canonical alpha cells, beta-associated ambient scores against canonical beta cells, and ribosomal and mitochondrial transcript fractions against canonical endocrine populations excluding αβδ cells. Donors with fewer than 10 cells in either comparison group were excluded from donor-level analyses.

### Disease Dataset Collection

Single-cell and single-nucleus RNA sequencing datasets from pancreatic disease conditions, including non-diabetic controls, type 1 diabetes, type 2 diabetes, pancreatic ductal adenocarcinoma, were collected from publicly available repositories. Datasets were selected based on data quality, availability of metadata, and compatibility with downstream integration. Where necessary, relevant metadata, including donor identity, disease status, and technical variables, were curated and standardized across studies to enable comparative analyses.

### Healthy and disease extension

Single-cell and single-nucleus RNA sequencing datasets spanning non-diabetic tissue (ND) and five pancreatic disease conditions (autoantibody-positive (AAB), type 1 diabetes (T1D), type 2 diabetes (T2D), and pancreatic ductal adenocarcinoma (PDAC)) were collected from publicly available repositories^6,9,68–72,97^ (Supplementary Table 1). Pre-aligned count matrices were quality-controlled per dataset independently using MAD-based outlier detection on log-transformed total counts, gene counts, and top-20-gene count fractions (threshold: 5 MADs), with additional filtering of cells exceeding 3 MADs above the median mitochondrial fraction. Doublets were identified and removed per sample using Scrublet, and genes expressed in fewer than 3 cells were excluded. Raw counts were retained for downstream analysis. Reference Mapping and Label Transfer: Extension datasets were mapped onto the core atlas using scANVI surgical mapping (scvi-tools v1.4.0), with a separate surgery model trained per condition. Extension cells were projected into the reference latent space using load_query_data with frozen dropout, treated as unlabelled during training, and with donor identity included as the batch covariate. Cell type labels were assigned at Level 4 resolution based on maximum posterior class probability, with coarser hierarchy levels derived deterministically by string parsing of Level 4 labels. Per-cell mapping confidence and normalised Shannon entropy were computed as QC metrics but were not used for filtering.

### Identification of diabetes-associated candidate cell states

Diabetes-associated candidate endocrine states were identified using uncertainty estimates from scANVI predictions and evaluated independently for type 1 diabetes (T1D), type 2 diabetes (T2D), and autoantibody-positive (AAB) datasets. Level-4 posterior probabilities were aggregated into Level-3 lineage categories by collapsing substates to their parent lineage. For each cell, lineage assignment confidence and normalized Shannon entropy were calculated from the resulting probability distributions. To identify disease-associated states while preserving endocrine identity, analyses were performed separately for alpha-, beta-, delta-, and pancreatic polypeptide (PP)-cell lineages within each disease context. Cells with strong lineage support (maximum Level-3 probability >0.8) were retained, and lineage-specific Level-4 probabilities were renormalized. A composite uncertainty score was calculated as the mean of normalized entropy, (1 − maximum posterior probability), and (1 − posterior probability margin between the highest and second-highest predictions). Candidate disease-associated cells were defined within each lineage and disease context as cells with composite uncertainty scores above the lineage-specific 75th percentile, while retaining strong lineage support (total lineage probability >0.75) and concordant hard lineage assignments. Candidate cells received disease-specific lineage annotations (for example, *Beta Cell_T2D*), whereas remaining cells retained their original annotations for downstream analyses.

### Distance calibration against healthy endocrine variation

To quantify the extent to which candidate disease-associated endocrine cells deviated from healthy endocrine variation, we calibrated cell-to-reference distances using held-out healthy donors. Healthy donors were randomly partitioned into reference and held-out subsets at the donor level. A k-nearest-neighbour (k = 50) graph was constructed using reference healthy endocrine cells in the scANVI latent space. For each held-out healthy cell and each candidate disease-associated endocrine cell, cosine distances to the 50 nearest healthy reference neighbours were calculated and summarized as the mean neighbour distance. The distribution of distances observed for held-out healthy cells was used to define the expected range of healthy endocrine variation. Disease-associated cells were then compared against this distribution, and the fractions exceeding the 95th and 99th percentiles of held-out healthy distances were calculated separately for each disease group and endocrine lineage.

### KNN Similarity

To quantify how disease-associated endocrine states related to healthy endocrine diversity, nearest-neighbour similarity was computed in the scANVI latent space using cosine distance. For this analysis, scANVI latent vectors were L2-normalized before nearest-neighbour search. Each query cell was assigned its k = 50 nearest healthy endocrine neighbours. Disease-associated states containing fewer than 10 cells were excluded. For each disease-associated state, neighbourhood composition was summarized as the fraction of nearest neighbours assigned to each healthy Level_4_extension_diabetes_simplified state. To provide a healthy comparator, the same procedure was applied to held-out healthy alpha and beta cells projected onto the remaining healthy reference cells. Neighbour-state compositions were visualized as heatmaps and used to quantify occupancy of healthy endocrine neighbourhoods. Per-cell neighbour assignments were retained, including cosine distance and cosine similarity, defined as 1 − cosine distance. To assess the proximity of disease-associated cells to healthy endocrine variation, disease-associated cell-to-reference distances were compared with distances obtained from held-out healthy endocrine cells projected onto the same healthy reference.

### Weighted Shannon Entropy

To quantify the specificity with which query cells mapped onto reference cellular states, we computed a normalized weighted Shannon entropy score based on k-nearest-neighbour relationships in L2-normalized latent embeddings using cosine distance. For each query cell, neighbouring reference cells were identified and weighted inversely by distance (epsilon = 10e-6). Weights were aggregated across reference state labels and normalized to generate a probability distribution across reference neighbourhoods. Shannon entropy (base 2) was calculated from this distribution and normalized by log2(K), where K represents the number of unique reference states, yielding scores ranging from 0–1. Lower values indicated mapping to a restricted set of reference states, whereas higher values reflected diffuse occupancy across multiple reference states. Dominant neighbour identities and their associated weights were additionally recorded for each cell. For diabetes analyses, candidate disease-associated alpha- and beta-cell states were projected onto healthy HPCA endocrine states using k = 50 neighbours in scANVI latent space. Entropy distributions and dominant-state frequencies were summarized separately for disease conditions, and pairwise comparisons were assessed using two-sided Mann–Whitney U tests. For stem-cell analyses, SC-islet cells were projected onto healthy HPCA endocrine states using k = 100 neighbours in scVI latent space to quantify the diversity of human endocrine-state occupancy.

### Integration of healthy exocrine epithelium and PDAC query mapping

Healthy acinar and ductal cells (Level 4) were extracted from the integrated human pancreas atlas. HVGs were selected globally and within each Level 3 exocrine lineage (Seurat v3, top 500 per selection), unioned with curated acinar and ductal state markers, and ambient-associated genes (exocrine digestive enzymes and islet hormones) were removed. Integration used scArches: SCVI was trained on raw counts (four hidden layers, 20 latent dimensions, batch and Level 4 labels encoded; up to 50 epochs with early stopping on validation ELBO), followed by SCANVI (40 epochs; unlabeled category, “Unknown”). Latent embeddings were visualized by UMAP (cosine neighbours, k = 50; min_dist = 0.75). PDAC acinar, ductal and malignant epithelial cells harmonized to the atlas gene space were mapped onto this reference with SCANVI.load_query_data() (dropout frozen). Missing reference genes were zero-padded. All PDAC cells were treated as unlabelled and the model was fine-tuned for 40 epochs (weight decay = 0). Joint latent space and SCANVI predictions were computed on concatenated healthy and PDAC cells, and UMAP was run on the combined embedding (cosine neighbours, k = 100; min_dist = 0.75). Analyses were performed in Python with Scanpy, scvi-tools/scArches, PyTorch and AnnData.

### Sample Representations from SampleCLR

To compare pancreatic donors across disease contexts while accounting for technical variability, we learned a 128-dimensional embedding for each donor–dataset sample by pooling scANVI embeddings with SampleCLR. Samples with fewer than 250 cells were removed; the remainder were split 80:20 for training and validation (seed = 0). Sample metadata (disease, age, sex, BMI, HbA1c, and technical covariates) were aggregated per donor with <u>patpy.pp</u>.extract_metadata. The model used a single-layer encoder (width 128) and a four-head attention aggregator (two layers, dropout 0.2) trained with the XSampleCLR contrastive loss (τ = 0.1; batch size 8; 250–500 cells per sample per step). A Batch-aware sampler (batch key Dataset, Pd = 0.9) reduced study-of-origin confounding. Optimization used Adam (learning rate 3 × 10⁻⁴, weight decay 1 × 10⁻⁴) with early stopping based on validation loss. Training comprised (1) 200 epochs of self-supervised contrastive learning and (2) up to 300 epochs of fine-tuning with a disease classifier (healthy, primary tumours, metastatic lesions, PanIN), weighting the contrastive loss at 0.1 relative to classification. We built a sample-level AnnData object (PDAC: 280 donors) storing fine-tuned (X_sampleclr_ft) and self-supervised (X_sampleclr_ssl) coordinates, cell-state composition percentages across Level_4 labels, and harmonized clinical metadata for downstream trajectory and pseudobulk analyses.

### Diffusion map embedding and diffusion pseudotime inference

On the sample-level SampleCLR graph, we built a k-nearest-neighbour graph from fine-tuned embeddings (X_sampleclr_ft; k = 20) and computed a diffusion map with Scanpy (sc.tl.diffmap, neighbours_key = neighbours_sampleclr_ft). Samples were visualized using selected diffusion components. Pseudotime was inferred with diffusion pseudotime and run on SampleCLR distances (distances_key = distances_sampleclr_ft). The root was set to a healthy reference donor for PDAC analyses by selecting the donor with highest composition homeostatic cell types. Sample-level object was used for downstream statistics and donor-level pseudobulk analyses.

### Pseudobulk gene–trajectory association analysis

To identify transcriptional programs associated with progression along donor-level trajectories, lineage-specific pseudobulk profiles were generated from cells belonging to the relevant cellular compartments. Raw counts were aggregated per donor, normalized to a library size of 10^6, log-transformed, and filtered to remove mitochondrial, ribosomal, hemoglobin and other lncRNA related genes. Genes detected in fewer than five pseudobulks were excluded. Pseudobulk samples were annotated using donor-level metadata including diffusion pseudotime, age category, suspension type and disease status. For each gene, associations between pseudobulk expression and diffusion pseudotime were tested using ordinary least squares regression while adjusting for suspension type:

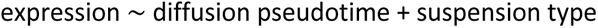

Categorical covariates were one-hot encoded using drop-first coding, and samples with missing covariate information were excluded on a per-model basis. Regression coefficients and associated P values for the pseudotime term were extracted for each gene and corrected for multiple testing using the Benjamini–Hochberg procedure. For analyses involving bifurcating trajectories, pseudotime–expression associations were additionally evaluated independently within trajectory branches by restricting models to donor subsets corresponding to each branch. Genes passing significance and effect-size thresholds were retained for downstream comparison of shared and branch-specific transcriptional programs.

### Gene-expression dynamics along pseudotime

To visualize transcriptional changes along the epithelial remodeling trajectory, diffusion pseudotime values were calculated from the donor-level manifold as described above. For each gene of interest, normalized single-cell expression values were modeled as a smooth function of pseudotime using generalized additive models (GAMs) implemented in the pyGAM package. Models were fitted using cubic spline basis functions with pseudotime as the predictor variable. Predicted expression values were evaluated on a uniformly spaced pseudotime grid spanning the observed range of the trajectory. Smoothed expression curves were plotted together with 95% confidence intervals derived from the fitted GAM. This approach was used to summarize gene-expression dynamics while accommodating non-linear transcriptional changes along the remodeling trajectory.

### PyDESeq2 endpoint differential expression

Pairwise contrasts between two endpoint states were tested with PyDESeq2 (v0.5.4). For each contrast, pseudobulks were restricted to the reference and test groups; at least two samples per group were required (min_samples_per_group = 2). Integer raw counts from layers[’counts’] were supplied to DeseqDataSet. Covariates were included only if present, complete, and variable (e.g. suspension_type when both nucleus and cell suspensions were represented). The design was additive: covariates plus the group factor, with covariates listed before the group term (design_factors = covariates + [group_col]). The reference level was set explicitly (ref_level = [group_col, ref]). dds.deseq2() was run, then DeseqStats with contrast [group_col, test, ref]. Results were sorted by adjusted P value and log2 fold change. Differentially expressed genes were defined as Benjamini–Hochberg padj < 0.05.

### eQTL-GWAS colocalization data

Pre-computed eQTL-GWAS colocalization results were obtained from Piron et al. 2025^130^, who applied the colocRedRibbon pipeline (direction-of-effect prefiltering and RedRibbon overlap followed by coloc.abf) to bulk pancreatic islet eQTLs from the TIGER^131^ consortium against T1D and T2D GWAS summary statistics^132–135^. We retained locus–gene pairs with PP.H4 ≥ 0.8. Gene symbols, lead SNP rsids, eQTL effect direction, eQTL beta, and SNP_PP_H4 were taken from Table S2 of the source publication. We utilised cellink^136^ for downloading, preprocessing and management of the summary statistics.

### Disease-state enrichment and eGenes in HPCA atlas sub-state expression mapping

Raw count matrices were normalised to 10,000 counts per cell and log(x + 1)-transformed. Mean log-normalised expression was computed per cell sub-state per condition using scanpy groupby aggregation, restricted to sub-states with ≥ 20 cells and ≥ 2 donors per condition. Sub-states were ranked by mean expression within the canonical eQTL cell type. This mapping uses expression data only and does not constitute a new eQTL analysis. Donor-level log2FC in cell-state proportion was computed using per-donor proportions as the statistical unit, with Mann–Whitney U tests and Benjamini–Hochberg FDR correction. Per-sub-state expression log2FC was computed as log2((mean_disease + 1×10⁻⁴) / (mean_healthy + 1×10⁻⁴)), where means were taken across all cells in that sub-state per condition group. For each colocalized eGene for which the IRA (increasing-risk allele) direction field was reported in Table S2 of Piron et al., concordance was defined as: concordant if eqtl_direction = ’inc’ and expression log2FC (disease/healthy) > 0 in the peak sub-state, or eqtl_direction = ’dec’ and log2FC < 0; discordant otherwise. This analysis was restricted to T2D eGenes. T1D sub-state expression comparisons are severely underpowered: end-stage T1D donors retain few beta cells such that per-sub-state means are dominated by noise rather than biology. To evaluate the contribution of T2D-associated cell-state loss to apparent discordance, concordance was stratified by the abundance log2FC of the peak sub-state derived from donor-level proportion comparisons. Sub-states with abundance log2FC < −1 showed only 37–41% concordance, consistent with expression comparisons being confounded by residual, biologically distinct surviving cells. Peak sub-states were therefore classified as compositionally stable (|abundance log2FC| < 0.5), depleted (log2FC < −0.5), or expanding (log2FC > 0.5), and concordance rates compared across categories. Peak sub-states were therefore classified as compositionally stable (|abundance log2FC| < 0.5), depleted (log2FC < −0.5), or expanding (log2FC > 0.5).

Highlighted eGenes were selected on two orthogonal, pre-specified criteria. The first criterion was breadth of sub-state concordance, the fraction of all valid sub-states showing directionally concordant expression. The overall top-ranked T2D eGene by this metric was MTNR1B (rs10830963; PP.H4 = 1.00), encoding the melatonin receptor and a well-established T2D risk gene, which was concordant across all 17 of its detectable sub-states. PHB (rank 5 of 185; majority_frac = 1.0 across all 73 detectable sub-states) was selected as the top biologically interpretable T2D eGene by breadth-of-concordance; its role in mitochondrial function and the T2D remodeling trajectory is discussed elsewhere in the manuscript. The second criterion was genetic-support strength, PP.H4, restricted to eGenes with concordant expression in a compositionally stable peak sub-state (stable_peak_concordant = 1.0; 93 of 185 eligible). FXYD2 (rs529623; PP.H4 = 0.999) ranked third of these 93 eGenes and was selected as the highest-PP.H4 T2D beta-cell eGene: the two genes ranked above it, MTNR1B (already captured by the first criterion) and ADCY5 (PP.H4 = 1.00; peak sub-state: Ductal Cell: Injury Reactive), are either redundant or peak outside the endocrine compartment. Both MTNR1B and ADCY5 additionally showed broad sub-state concordance (majority_frac = 1.00 and 0.69 respectively), consistent with the finding that high-confidence T2D loci tend to show directionally concordant expression changes across multiple pancreatic cell states.

### TCGA Survival Analysis

TCGA-PAAD^137^ bulk RNA-seq expression and clinical survival annotations were used to assess whether epithelial remodeling programs identified in the atlas were associated with patient outcome. Gene-level TPM values were log-transformed as log₂(TPM + 1) and merged with patient-level clinical data. Overall survival time was defined as days to death when available, otherwise days to last follow-up; event status was defined from recorded vital status (dead = event). A malignant remodeling program score was calculated as the mean z-scored expression across patients of genes in the malignant remodeling program (S100P, TMPRSS4, MSLN, PSCA, MUC5AC, GPRC5A, KRT6A, SEMA3C, TNS4, CLIC3, AHNAK2, and ONECUT3). For individual-gene analyses, log₂-transformed expression of SEMA3C, TNS4, CLIC3, and AHNAK2 was used directly. Patients were stratified using the upper and lower quartiles of each score or gene, excluding patients with intermediate values. Kaplan–Meier survival curves were estimated for the upper- and lower-quartile groups, and survival differences were assessed using two-sided log-rank tests.

### Presence Score

Similar to HNOCA (He et al. 2024)^118^, we quantified how strongly each human endocrine cell in the pancreas atlas is represented by mouse diabetes models using a weighted cross-species nearest-neighbour framework followed by diffusion smoothing, yielding a per-cell presence score for each model (e.g. NOD, db/db, mSTZ-based models). Integrated latent embeddings were used for cross-species comparison (human scVI embedding; mouse integrated embedding). Analysis was restricted to matched endocrine compartments (human Level 4 endocrine states; corresponding mouse endocrine cell types). For this compartment, we built a mouse-to-human k-nearest-neighbour graph with NNDescent (k = 100), using bidirectional query–reference neighbour sets to define edges. Edge weights reflected local neighbourhood concordance via overlap of human kNN sets (k = 100): the number of shared neighbours between a mouse and human cell was converted to a Jaccard index and squared. Mouse cells were aggregated by diabetes_model: for each model, row-normalized edge weights were summed across that model’s mouse cells and divided by model size to obtain an initial coverage vector over human cells. To denoise and propagate signal across nearby human cells, we constructed a human-only kNN graph (NNDescent, k = 50), symmetrized it, and derived a transition matrix for random walk with restart (RWR; α = 0.5, 100 iterations). Resulting vectors were transformed with log1p, winsorized to the 5th–95th percentiles, and min–max scaled to [0, 1] per model to obtain final presence scores.

### Comparison of mouse-model and human disease-context program distributions

HPCA endocrine states were collapsed into broad alpha- and beta-cell programs and low-entropy states were excluded. For each mouse model, normalized program-occupancy profiles were computed from mean neighbourhood presence scores across collapsed programs. Human disease-context distributions were calculated as the fraction of endocrine cells assigned to each collapsed program within healthy-control, T1D, T2D and autoantibody-positive cohorts. Similarity between mouse-model occupancy profiles and human disease-context distributions was quantified using cosine similarity across matched endocrine programs.

### Graph-propagated beta-cell program scoring

Program scores for SC-islet beta cells were derived using the integrated neighbour graph linking SC-islet query cells and core human beta-cell references. Query-to-reference connectivity weights were extracted and row-normalized, and for each query cell the fraction of connectivity assigned to predefined human beta-state groups was calculated. Human beta-state groups included mature identity, dedifferentiation, immediate-early activation, stress-associated states, proliferation and oxidative phosphorylation/metabolic states. Composite remodeling scores were calculated as the sum of dedifferentiation, activation, stress and proliferation programs, and maturity–remodeling balance was defined as mature identity minus remodeling/plasticity scores. Program scores were summarized across maturation stages for downstream analyses.

### Adult beta-cell attractor distance and fidelity

An adult beta-cell attractor centroid was defined as the coordinate-wise median scANVI embedding of mature core human beta cells corresponding to INS-high, mature identity and secretory granule states. For each beta cell, cosine distance to the attractor was calculated and fidelity was defined as one minus the attractor distance. Distances and fidelity values were computed for human and SC-islet beta cells and summarized across maturation stages. Neighbour graphs, UMAP embeddings and diffusion maps were generated on the combined human and SC-islet beta-cell subset using scVI latent representations.

### Identification of attractor-associated genes

Analyses were performed in core human beta cells following library-size normalization, log transformation and filtering to remove mitochondrial, ribosomal and hemoglobin-associated genes. Genes detected in fewer than five cells or expressed in fewer than 5% of cells were excluded. Associations between gene expression and beta-cell fidelity scores were quantified using Spearman correlation. P values were adjusted using the Benjamini–Hochberg procedure. Genes passing predefined significance and effect-size thresholds were retained for downstream analyses.

### Gene signature analysis across SC-islet maturation

Curated attractor-associated and peripheral gene signatures were scored in normalized SC-islet beta cells using Scanpy gene-set scoring. Signature scores were summarized across maturation stages together with attractor distance and fidelity metrics. Comparisons between early-stage, late-stage and mature human beta-cell expression profiles were performed for shared genes, and convergence ratios were calculated relative to mature human beta-cell reference expression. Selected genes were visualized using z-score normalized expression values across maturation stages.

## Code and data availability

The datasets can be downloaded from public sources (Supplementary Table 2). The processed Atlas will be available on CAP and the code will be made public upon publication.

## Acknowledgments

This project was supported in part by grant number CZIF2022-007488 from the Chan Zuckerberg Initiative Foundation. S.P and L.A. were supported by the Helmholtz Association, as part of the joint research school Munich School for Data Science (MUDS). This work was supported by the Deutsche Forschungsgemeinschaft (DFG; project number 515571394, grant number TH 900/18-1; S.P.); the Chan Zuckerberg Initiative Foundation (CZIF; grant CZIF2022-007488, Human Cell Atlas Data Ecosystem; S.P., M.D.L.); the DFG Leibniz Prize awarded to F.J.T. (J.L.B.); the European Union (ERC, DeepCell, grant 101054957; F.J.T., E.R.); the Studienstiftung des deutschen Volkes (L.A.); the Innovative Medicines Initiative 2 Joint Undertaking (JU; grant agreement No. 101007873, EU-IMI2 101007873; E.R.); and Breakthrough T1D (grant number 1-INO-2024-1545-A-N; S.J.).

## Author contributions

S.P., D.C.S., F.J.T., M.D.L. and D.B. conceived the study. S.P. and D.C.S. contributed equally and have the right to list their name first in their curriculum vitae.

S.P., and D.C.S. led, and organized the project with support from S.J. S.P. and D.C.S. led the analysis of the atlas, with support from S.J, J.L.B, L.A, E.R.

M.D.L, D.B. and F.J.T. supervised the work.

All authors from the HCA Pancreas Bionetwork contributed to annotating the atlas and guided the analysis.

F.J.T. acquired funding. All authors contributed to writing, reviewing and editing the manuscript, and approved the final version.

## Supplements

Supplementary figures HPCA

[HPCA ATLAS SUPPL TABLE](https://docs.google.com/spreadsheets/d/1EmuCpLQexWK2f_zJkR4O1SQPcVrdlrl1xciTL8BpUL8/edit?gid=0#gid=0)

